# IRIS: A Machine Learning-Based Pose Re-Ranking Tool for RNA-Ligand Docking

**DOI:** 10.1101/2025.10.24.684232

**Authors:** Andrew Amburn, Shalini J. Rukmani, Jerry M. Parks, Jeremy C. Smith

## Abstract

RNA-ligand docking remains challenging, due in part to intrinsic properties of RNA such as structural flexibility and a highly charged phosphate backbone. rDock, a widely used RNA docking program, can generate ligand poses close to the experimental structure, but its scoring function frequently fails to rank these poses above less accurate alternatives. To supplement rDock, here we introduce the Intelligent RNA Interaction Scorer (IRIS), a regression model leveraging physicochemical and interaction-based features and trained on the largest dataset of experimental nucleic acid-ligand complexes compiled to date (1,356 structures). IRIS improves rDock RNA-ligand pose ranking relative to the use of rDock scores alone. We find that at least one of the 100 top generated poses for any given complex is within 2.0 Å RMSD of the native pose in 79.4% of test complexes. Of these 79.4%%, the default rDock scoring function ranks the correct pose first in 40.2% of cases. IRIS improves this latter fraction to 52.7% and increases the success rate for selecting a near-native pose among the top five ranked poses from 55.4% to 73.2%. IRIS thus significantly enhances pose ranking accuracy and can be seamlessly integrated into docking pipelines to refine ligand poses in RNA-targeted drug discovery.

## Introduction

RNA plays diverse roles in cellular regulation and catalysis, functioning as riboswitches, ribozymes, and scaffolds for macromolecular assemblies^1^, and have also been implicated in a wide range of human diseases, including cancers and infectious, immune, cardiovascular, and neurodegenerative disorders^2–6^. RNA targeting thus offers a promising route to expand the druggable genome space and can include genes that are non-coding or prove challenging to target at the protein level^7^. RNA molecules are thus attractive targets for drug discovery campaigns, and RNA targeting has gained significant momentum over the last decade^8,9^.

Computational docking is a promising approach to identifying high-affinity small-molecule inhibitors for macromolecular targets. Docking programs can predict the preferred 3D orientation and conformation of a ligand bound to a given receptor (referred to as the pose) with an estimated receptor-ligand binding score or affinity^10^. These programs use physics-based, knowledge-based, ML-based, or hybrid scoring functions to identify the most probable binding mode of a ligand to a receptor of interest. The accuracy of the scoring functions, however, remains a primary limitation in molecular docking applications, and especially for RNA systems^11–13^. In contrast to protein-targeted drug discovery, which has seen notable success with docking^14–17^, RNA-targeted screening presents unique challenges^1,7^. RNA molecules contain highly charged backbones with phosphate groups, leading to the participation of coordinating metal ions and water molecules to mediate ligand-receptor interactions and stabilize the binding pocket and tertiary structure folds^18–21^. Also, RNA molecules can adopt complex tertiary structures specific to their function (that include, but are not limited to, kissing loops, pseudoknots, G-quadruplexes and coaxial stacking) with molecular driving forces including electrostatic stabilization by water and metal ions and base pair stacking. Further, the structural changes induced by the presence of ligands vary considerably between different RNA systems, from highly specific conformational transitions (on/off state) in certain riboswitches (e.g. SAM-I) while shifting equilibrium towards a given conformation from multiple possible conformations in some RNA systems (e.g. Group-II intron)^22^. Capturing the RNA-small molecule interactions in highly polar, flexible and low buried surface area RNA binding sites is thus challenging^23–26^. Further, generating representative conformations of RNA-binding small molecules (including protonation states and tautomers) requires robust approaches as these molecules may exhibit physicochemical properties distinct from protein small-molecule inhibitors, including lower octanol-water partition coefficients (LogP), higher topological polar surface areas (TPSA), higher numbers of hydrogen bond donors and acceptors, and higher numbers of heteroatom-containing rings [1]. Finally, a significant other challenge in RNA docking is the low number of available experimental PDB structures compared to proteins^13,27^.

Comparative assessments have been performed on a wide range of docking programs, including those originally developed for protein–ligand interactions, to evaluate their effectiveness on nucleic acid–ligand complexes^11,27^. These studies reveal that performance varies widely depending on the docking protocol and target type, and they underscore the need for tools that perform reliably in RNA-ligand docking scenarios. Among the docking programs evaluated, rDock has emerged as perhaps one of the more accurate programs for RNA-ligand pose generation and one of the few that can perform both local docking (in which the binding site is known) and global docking (in which the binding site is unknown) with reasonable accuracy. rDock has demonstrated one of the most competitive capabilities for predicting near-native conformations given sufficient sampling but arguably suffers from the lack of a scoring function that can consistently rank such poses correctly^11^.

In addition to the standalone docking programs, alternate scoring functions that use knowledge-based, ML-based, or a hybrid of these two have been developed over the years to improve pose prediction accuracy^28–30^ (e.g. RNAPosers, AnnapuRNA, and SPRank). Several of these scoring functions use rDock to generate the ligand-RNA poses for refinement and re-ranking. Further, direct scoring methods using data-driven strategies are being increasingly pursued to circumvent expensive docking runs but so far have not seen high success rates and are yet to be tested on a large-scale^7^.

A common challenge historically faced by many RNA specific scoring functions is the lack of solved experimental structures of RNA-ligand complexes, both in terms of quantity and quality to be used for validation^28–30^. The development of new, improved scoring functions and programs building upon the existing data is thus a cumbersome process but can be accelerated greatly by ML methods. A recent effective example of an RNA-specific ML scoring function, to which we incorporate a comparison in the present work, is RNAPosers, which was developed to improve pose ranking for RNA–ligand complexes using classifiers trained on experimental structures^29^. RNAPosers represents each ligand pose using a high-dimensional “pose fingerprint” that encodes distances between all heavy atoms in the ligand and all nearby heavy atoms in the RNA receptor. With a leave-one-out training and testing approach, these classifier models recovered poses that were within 2.5 Å of the native poses in ∼80% of the 80 cases examined, and, on two separate validation sets, recovered such poses in ∼60% of the cases.

The performance of ML-based methods is heavily dependent on the quality and quantity of training data. Insufficient or biased training sets can lead to inaccurate predictions and ML models perform best when there is a large, diverse set of high-quality training data available. Selecting the appropriate features to represent the receptor-ligand interactions is also extremely important, and improper feature selection can lead to poor model performance^28^. Most previous models have been trained on interaction data extracted from experimentally solved complexes obtained from the Protein Data Bank (PDB)^31^, the Nucleic Acid Knowledgebase (NAKB)^32^, or from the literature, and have used specific sets of docking software sub-scores and 2D and 3D physicochemical descriptors of the ligand, the RNA binding site, and interactions between the RNA and ligand^29,33^. However, to our knowledge, most current ML-based scoring functions for RNA have not been trained on the full set of available ligand-bound structures and do not include an optimal number of features with respect to their training and validation data sets, or both. Furthermore, although DNA-ligand complexes are highly similar in structure to RNA-ligand complexes, most, if not all current RNA-based ML scoring functions do not include DNA-ligand complexes in their training sets.

To address the limitations present in current ML-based RNA-specific scoring functions, we have designed and tested an ML-based re-ranking tool for RNA docking that we call the Intelligent RNA Interaction Scorer (IRIS). This method uses a collection of DNA and RNA-ligand complexes in its training set. rDock was used to generate the ligand-RNA poses considering its consistent performance in being able to generate near-native poses as mentioned earlier. IRIS uses selected rDock sub-scores together with diverse 2D and 3D physicochemical descriptors as features. We compare the results against the rDock default scoring functions for global and local docking as well as RNAposers. Our results emphasize the potential of approaches such as IRIS as effective predictive tools for RNA-small molecule ligand design. IRIS is freely available at https://github.com/AndrewAmburn/IRIS.

## Material and Methods

### Dataset compilation

Our dataset includes 1,326 structures of nucleic acid (NA)-ligand complexes taken from the Protein Data Bank (PDB RCSB)^31^ and the Nucleic Acid Knowledgebase (NAKB)^32^. Given this rather limited number of available NA–ligand complexes, to maximize coverage we did not apply filters based on resolution, experimental method (e.g., X-ray, NMR, cryo-EM), sequence length, or receptor type. All entries containing both a nucleic acid chain and at least one bound small-molecule ligand were included. For complexes with multiple unique ligands bound to a single receptor, each ligand–receptor pair was treated as a separate complex. For each such complex, any co-crystallized ligands other than the ligand designated for re-docking were removed prior to docking to simulate the docking of ligands to NA-receptors in which additional bound ligands are unknown, as is the case in many HTVS screening campaigns. This expanded the original dataset to 1,435 3D structures of NA-ligand complexes. To our knowledge, this dataset, available in the Supplementary Information, is the largest to date for any ML-based tool designed for RNA docking.

### Redundancy filtering

To prevent information leakage between training and test sets, we applied stringent redundancy filtering based on both ligand identity and receptor sequence similarity. Specifically, a complex was considered redundant only if it shared an identical ligand (as defined by canonical isomeric SMILES) and ≥90% sequence identity in its nucleic acid receptor with another complex in the dataset. Receptor sequences were extracted from their respective PDB files by mapping nucleotide residue names to one-letter base codes (A, C, G, U, T). Sequence identity between receptors was computed using Biopython’s^34^ global pairwise alignment without gap penalties (globalxx). Ligand identities were determined using canonical isomeric SMILES, generated with RDKit^35^ to ensure consistent encoding of molecular structure and stereochemistry. The redundancy removal process resulted in 977 unique NA–ligand complexes.

This dual-filtering approach ensured that complexes with the same ligands but different receptor contexts, or complexes with the same or similar receptor but a different ligand, were retained, allowing for reduction of bias in testing and training sets and capturing of broader chemical and structural diversity across the receptor-ligand complexes. The resulting set of non-redundant complexes represent the full dataset we used to train and test our IRIS model. A complete list of redundant complex pairs and a per-complex summary of redundancy are provided in the Supplementary Information.

### Feature set selection

To comprehensively characterize each nucleic acid–ligand complex, we generated an initial set of 314 features that combined both receptor-, ligand-, and interaction-level descriptors. These included standard 2D and 3D molecular descriptors computed using RDKit ^35^, as well as custom features derived using PyMOL^36^, SciPy^37^, and in-house scripts that quantified electrostatic interactions, steric complementarity, spatial geometry, nucleobase composition, and distance-based metrics. Additional pose-specific features were extracted from the rDock docking output files, including energy-based subscores from the rDock scoring function (Equation 4), which capture distinct terms unique to each docked pose.

To reduce dimensionality and prevent multicollinearity, we eliminated features with zero, missing, or constant values across all complexes. We then applied pairwise Pearson correlation filtering to remove highly collinear features, defined as having an absolute Pearson correlation coefficient greater than or equal to 0.8. After filtering, a final set of 178 features was retained for model training and testing. The full feature set used in the IRIS model is available in the Supplementary Information.

### Train/test splitting

To construct a distinct and representative test set for model evaluation, we applied a 15% stratified sampling approach over the full set of 977 non-redundant NA–ligand complexes. For each complex, we first applied our feature generation pipeline to the native ligand pose. Then, following the iSIM method^38^, for each NA–ligand complex, we concatenate the feature set of molecular descriptors column-wise into a unified high-dimensional feature vector. Pairwise similarities were then computed using the Jaccard/Tanimoto (JT) coefficient^39^, defined as:

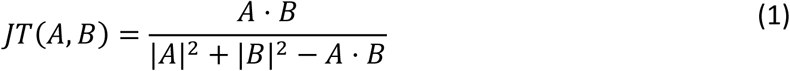

where *A* and *B* are continuous feature vectors for two complexes, and *A⋅B* denotes their dot product. For each complex, we calculated its average JT similarity to all others to obtain a complementarity similarity score (*comp_sim*). Complexes were then sorted in ascending order of *comp_sim*, and the sorted list was partitioned into n = ⌊ [0.15] × N⌋ strata. Although the model is designed for RNA–ligand pose re-ranking, DNA–ligand complexes are included in the training set. Thus, to evaluate whether the model generalizes beyond RNA, we intentionally included approximately 25% DNA complexes in the test set to assess whether IRIS could also improve pose ranking accuracy over rDock for DNA targets. This initial test set comprised 146 complexes, including 110 RNA and 36 DNA complexes, preserving an approximate 75:25 RNA:DNA ratio.

Although some chemical similarity among ligands is expected within the training set due to limited chemical diversity in publicly available NA-ligand complexes, we sought to ensure that our model did not contain large clusters of ligand chemotypes which may bias model training. To assess this, we clustered all training ligands based on Tanimoto similarity (≥0.8) using Morgan fingerprints^40^ (radius = 2, 2048 bits) from the RDKit library^35^. We then calculated the fractional representation of each ligand chemotype cluster relative to the full training set. The most frequent cluster accounted for only 4.07% of the total training data. This cluster contained almost exclusively spermine and glycerol ligands which are involved in stabilizing nucleic acids during structure resolution^41,42^. The top 10 ligand clusters collectively represented just 15.91% (**Table S1**) of the total training set. These results confirm that the model is likely not biased toward any dominant scaffold or ligand chemotype and that the training set exhibits substantial chemical diversity. A full description of each cluster is presented in the Supplementary Information.

To ensure that the test set was structurally independent from the training set, we performed a ligand-level similarity analysis. For each test ligand, we identified all training ligands with a Tanimoto similarity greater than 0.8, computed using Morgan fingerprints (radius = 2, 2048 bits). For each high-similarity pair, we computed the root-mean-square deviation (RMSD) between their maximum common substructures (MCS) using RDKit ^35^, without any structural alignment or minimization.

This analysis revealed five test ligands with an MCS RMSD ≤ 5 Å to a training ligand, indicating potentially near-duplicate poses. These complexes were removed from the test set to eliminate possible structural similarity to ligands seen in the training set (**Table S2**). The final test set consists of 141 NA–ligand complexes, comprised of 108 RNA and 33 DNA complexes.

To assess the chemical diversity of the selected test set, we applied t-distributed stochastic neighbor embedding (t-SNE)^43^ to project the complete feature set fingerprint matrix into two dimensions for each complex in the dataset (**Figure 1**). This non-linear dimensionality reduction method transforms high-dimensional data into a lower-dimensional representation by preserving local similarities and minimizing the Kullback–Leibler divergence^44^ between the high- and low-dimensional distributions. This objective function is represented as:

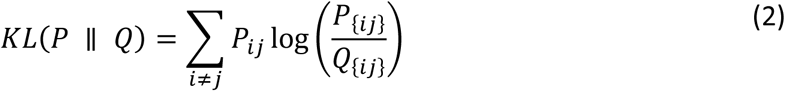

where, *P_ij_* and *Q_ij_* denote the pairwise similarity probabilities in the high- and low-dimensional spaces, respectively. The resulting projection coordinates, labeled t-SNE1 and t-SNE2, are abstract and do not correspond to original input features, but they reflect the local structure of the feature space. The test set compounds (shown in blue) are broadly distributed across the full t-SNE embedding of the dataset (shown in gray), indicating that the stratified sampling strategy effectively captured a representative range of chemical diversity within the NA–ligand interaction space.

**Figure 1.**
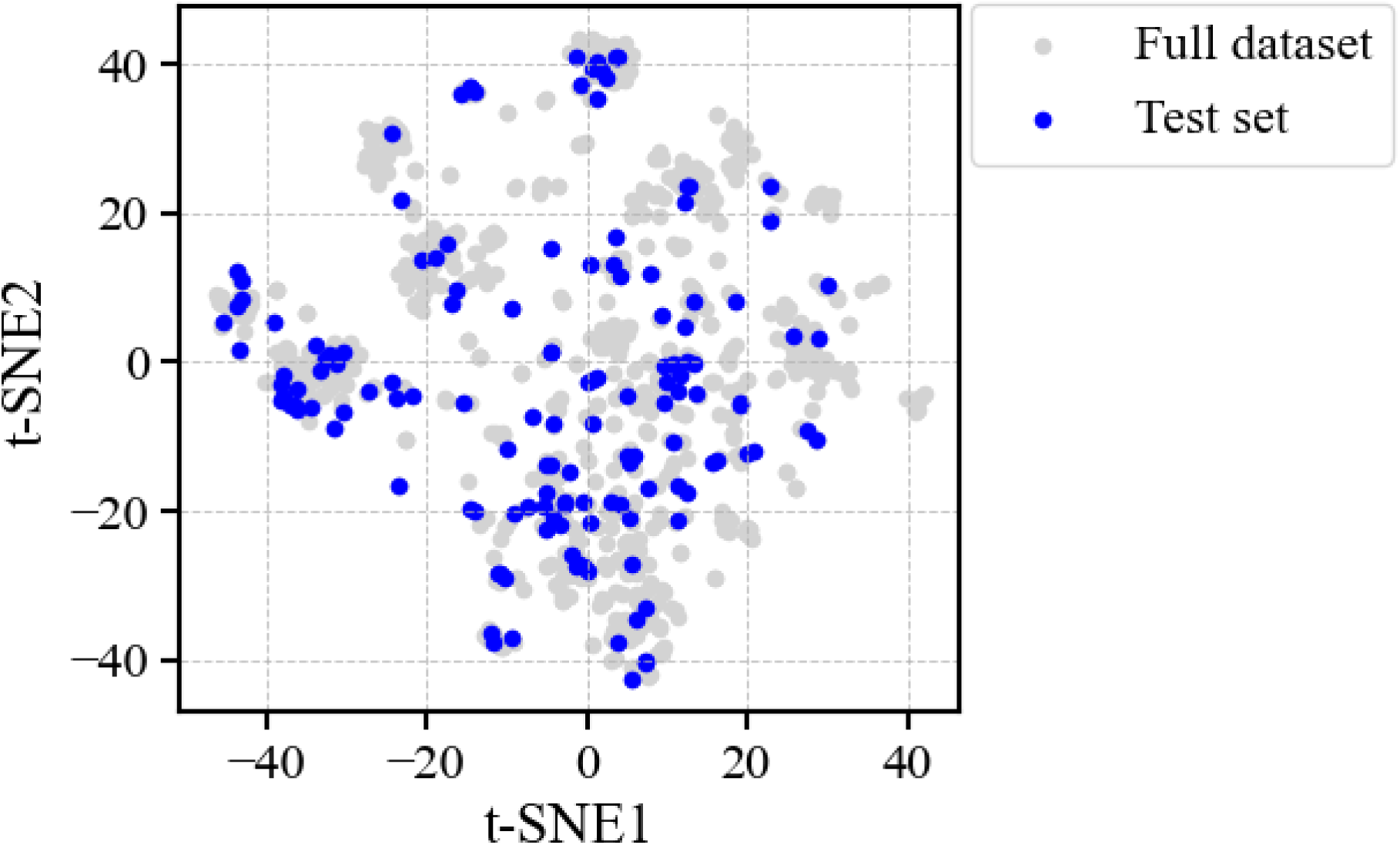
Distribution of stratified test set complexes in the t-distributed stochastic neighbor embedding (t-SNE) of the IRIS dataset. High-dimensional feature vectors for all 977 nucleic acid–ligand complexes were reduced to two dimensions using t-SNE. Gray points represent the full dataset, and blue points indicate the 141 held-out test complexes selected via complementarity-based stratified sampling using the Jaccard/Tanimoto similarity coefficient. Test set points are broadly distributed across the full feature space, confirming that the sampling strategy captured representative regions of the dataset.

### Intra-test set similarity analysis

To quantify the chemical diversity of the test set a Tanimoto similarity matrix^56^ was generated using Morgan fingerprints^40^ from the RDKit library^35^. Morgan fingerprints encode molecular structures by capturing local atomic environments within a defined radius, making them well-suited for quantifying molecular similarity. Ligand structures obtained from Structure Data Files (SDF) associated with test set complexes were first converted into RDKit Mol objects to ensure proper molecular parsing. Each ligand was then transformed into a 2048-bit Morgan fingerprint using a radius of 2, which encodes atomic connectivity up to two bonds away from each atom. This fingerprinting method captures both local and extended molecular features. To assess pairwise molecular similarity, the binary Tanimoto similarity coefficient (𝑇) was computed for all ligand pairs in the test set and is defined as:

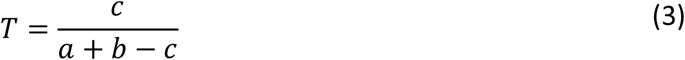

where *a* is the number of bits set to 1 in fingerprint A (training set), *b* is the number of bits set to 1 in fingerprint B (test set), and *c* is the number of bits set to 1 in both fingerprints A and B (the intersection of the two fingerprints). This metric quantifies the proportion of shared fingerprint features, with values ranging from 0 (no shared substructures) to 1 (identical molecular fingerprints). Importantly, the binary Tanimoto coefficient differs from the generalized Jaccard/Tanimoto formulation used earlier for continuous, high-dimensional descriptor vectors by being strictly defined on binary feature representations of molecular structure.

We then extracted any test set ligand complexes which exhibited a pairwise binary Tanimoto coefficient of 0.8 or higher (**Figure S1**). To evaluate whether these high-similarity pairs also exhibited spatial redundancy, we computed the root-mean-square deviation (RMSD) between their maximum common substructures (MCS) without alignment using RDKit^35^ (**Table S3**). Despite Tanimoto values of 0.8–1.0, all high-similarity ligand pairs had MCS RMSD values greater than 8 Å, and many exceeded 30–100 Å.

This result confirms that the selected test ligands are not only mostly chemically diverse but also occupy distinct conformational and geometric regions in 3D space if chemically similar. These findings validate that the test set is free of internal structural redundancy and provides a basis for evaluating pose re-ranking performance across a range of RNA- and DNA-ligand complexes.

### Ligand protonation and tautomer assignment

Assigning correct protonation and tautomeric states to small molecules in nucleic acid (NA) complexes remains a fundamentally unresolved challenge and is highly non-trivial for even a single NA-ligand complex. Considerations for nucleic acid environments are complicated by intense electrostatics, such as the presence of metal ions that may or may not coordinate ligand binding or stabilize conformational states^45^. In addition, structural data for NA–ligand complexes typically originate from crystallography, cryo-electron microscopy, or nuclear magnetic resonance (NMR) spectroscopy, where ligand confirmation, protonation and tautomeric states can be influenced by experimental conditions, such as nonphysiological pH, ionic strength, magnetic fields, and cryocooling rather than reflecting their native cellular state^46–48^.

Given these ambiguities, we adopted a standardized and widely used approach to hydrogen assignment by applying Open Babel’s command-line protonation routine^49^ (obabel -h) to all ligand and receptor structures. This tool adds explicit hydrogens so that each atom reaches its typical bonding capacity (“valence”) based on its element type and bonding geometry (“hybridization”), independent of explicit pH or local electrostatics. Notably, this same method is also used by other RNA-targeted machine learning models, including AnnapuRNA^28^, and enables consistent preprocessing across a large and chemically diverse dataset.

To maintain consistency with the experimentally determined structures, all ligands were retained in their original stereochemical and tautomeric forms as reported in the corresponding PDB entries. This approach avoids introducing artificial deviations from the native binding pose and aligns with practices used in prior RNA–ligand docking studies^50^.

### Treatment of structural metals, waters, and cofactors in docking

In our docking workflow, metal ions, structural water molecules, and bound cofactors present in the crystal structures were deliberately removed prior to pose generation. Although RNA-ligand binding is heavily influenced by metal- and water-mediated interactions, removal of these elements during docking is necessary and standard protocol in RNA-targeted virtual screening due to limitations in representation of these interactions by docking software. Specifically, rDock permits inclusion of these components but by default assigns metals a uniform formal charge of +1 for example, which do not accurately capture divalent metal coordination. In testing, the addition of explicit water molecules greatly increases computational expense for all complexes ^51^. Therefore, these components may result in vastly increased compute time and poor docking performance if included.^51^

Many recent RNA-ligand docking studies concur with this practice by discarding ions, cofactors, and structural waters to simplify the docking system^7,11,27,50,52^ and thereby avoid distortions due to poorly modeled electrostatics or poorly placed explicit solvent molecules.

Importantly, although these structural elements are omitted during docking, they are reintroduced during feature extraction for the IRIS model if they are known. Specifically, features which describe these interactions, such as the minimum distance from any atom in the ligand to any metal ion or water molecule coordinates, minimum distance from receptor to metal ions, are computed based on the coordinates of these structural elements in the experimental structure.

This ensures that the IRIS model incorporates the spatial context and potential interactions provided by these biologically relevant components if they are available. While our current approach follows established conventions, establishing a benchmark of the impact of retaining versus removing these structural elements during docking across diverse NA-ligand complexes would be beneficial for future studies to investigate.

### rDock docking methods

Ligand poses in rDock are generated through a multi-stage search process that combines stochastic and deterministic methods to explore the binding site^51^. For each ligand pose, this process begins with a series of three sequential genetic algorithm stages designed to diversify the search and prevent early convergence on suboptimal solutions. After the genetic algorithm stages converge on a single pose, pose refinement continues with a low-temperature Monte Carlo simulation, which introduces controlled perturbations to escape shallow local minima while preserving favorable interactions. This step is followed by simplex minimization, which locally optimizes the pose by adjusting atomic coordinates to reduce the total score^51^.

rDock provides two methods for molecular docking: the reference ligand (*RL*) method and the two sphere (*TS*) method. In the *RL* method, the binding site cavity is defined based on the coordinates of a known ligand from an experimental structure. A grid is placed around the ligand within a user-defined radius. Grid points that overlap with receptor atoms or fall outside the defined region are excluded. A small spherical probe (1.0–2.0 Å radius) is used to refine the cavity by excluding areas inaccessible to the probe, ensuring that only the search space surrounding the experimental ligand is considered during docking^51^. This method is suitable for redocking studies and cases in which the binding site is well characterized (i.e., “local docking”).

The TS method identifies potential binding cavities by placing a grid over a search space of a user-defined radius centered on specified coordinates within the receptor. The method uses two probe spheres of different user-defined sizes—a large sphere (3.5–6.0 Å radius) to exclude shallow, flat, or convex regions, and a small sphere (1.0–2.0 Å) to identify deep and accessible pockets suitable for ligand binding. Grid points accessible only to the small probe are retained, and user-defined thresholds such as minimum cavity volume (e.g., 100–300 Å³) control the size and shape of the cavity^51^. This method is useful for ‘global’ docking, i.e. when no prior knowledge of the binding site is available.

For both the *RL* and *TS* docking methods, in the present study 100 poses per complex were generated by invoking the *rbdock* engine with the *-n* 100 option. This performs 100 independent docking runs per ligand, in which each run begins with a randomized initial conformation within the identified binding site and optimizes the ligand’s position, orientation, and torsional angles using a Genetic Algorithm (GA). The GA iteratively evolves a population of candidate poses based on docking scores and selects the best-scoring pose from each run. As a result, 100 distinct docked poses are produced for each ligand-receptor complex, each representing a local minimum in *S_total_* identified from a different randomized starting point^12^. Importantly, since each of the 100 poses generated per complex by rDock is accompanied by a unique docking score, each of the 100 poses per complex differ in conformation to any other sampled pose. Detailed docking parameters for both *RL* and *TS* docking methods are given in the “rDock Docking Parameters” section of the SI.

### rDock scoring function terms

The generic rDock scoring function is a weighted sum of ligand and receptor intra- and inter-molecular interaction terms and is given by equation 1. *S_inter_* scores the intermolecular interactions between the ligand and receptor, *S_intra_* accounts for the relative intramolecular strain energy of the ligand, *S_site_* estimates the conformational flexibility of terminal OH and NH_3_ groups on the receptor within the binding site, and *S_restraint_* includes pharmacophoric restraint and cavity penalty terms by default and can also be used to bias poses based on prior knowledge [27].

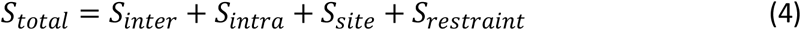

rDock uses two scoring functions, namely, *dock* (equation 4) and *dock_solv*. The latter scoring function includes a desolvation term (*S_solv_*) added to equation 1 to represent the change in solvation energy of the ligand and the receptor docking site upon ligand binding.

### IRIS model training workflow

The IRIS training workflow (**Figure 2**) begins with ligand-RNA docking using rDock, with pre-processed ligand SDF and corresponding receptor Mol2 files used as inputs to generate 100 poses per complex ranked by the default rDock scoring functions. Feature extraction was then performed to compute both ligand-level and pose-specific descriptors from the docking output, capturing chemical, geometric, and energetic properties of each ligand–receptor interaction. Ground truth labels were generated by calculating the root mean square deviation (RMSD) of each ligand pose relative to its experimental pose. Symmetry-corrected heavy-atom RMSDs were computed with the Python package *spyrmsd*^53^, which accounts for atomic equivalence and molecular symmetry. Docking poses were compared to the corresponding native poses after receptor alignment but without ligand superposition.

**Figure 2.**
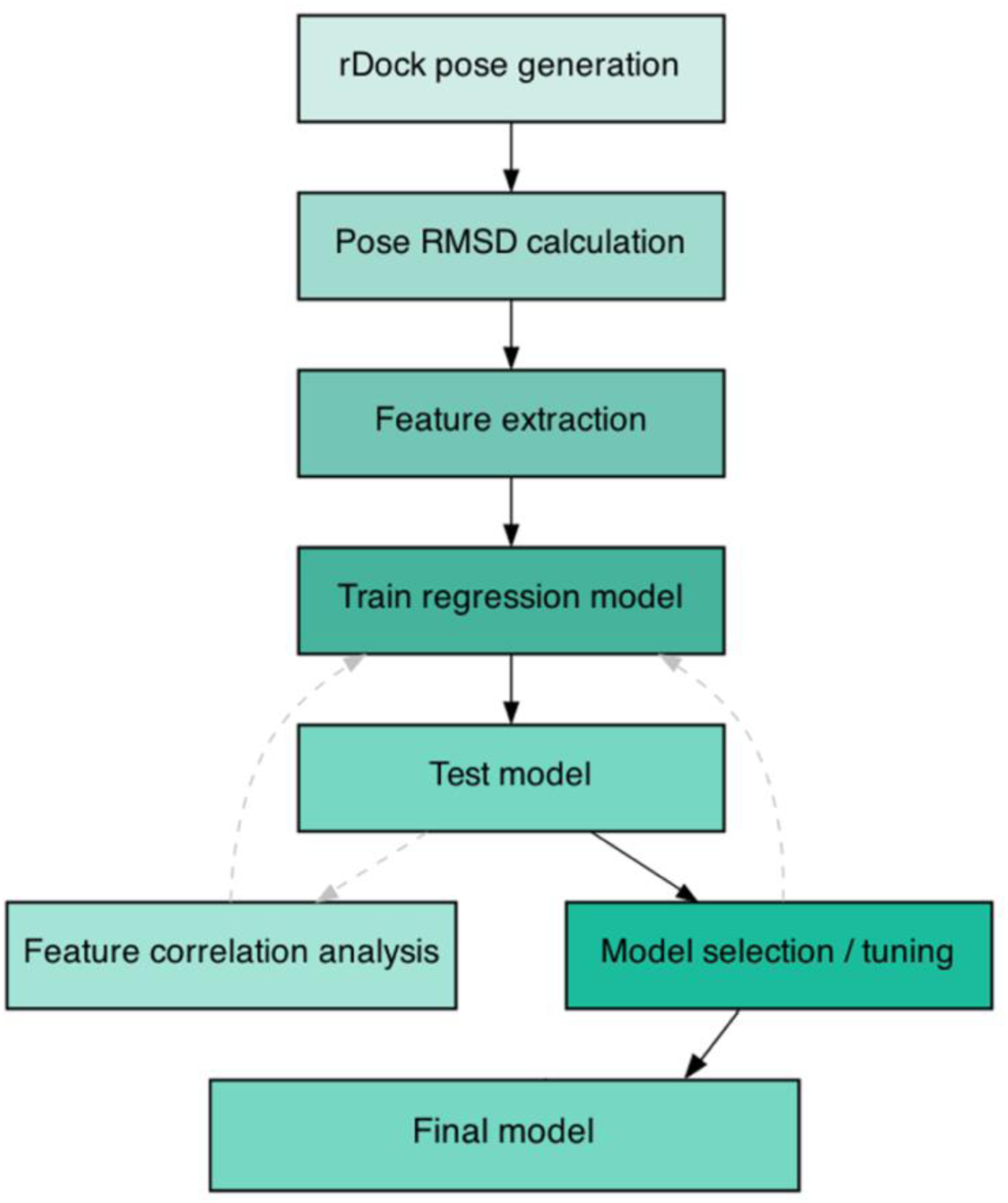
IRIS training workflow. Illustration of initial IRIS model generation and pose re-ranking validation on test set, including rDock runs to generate ligand poses, feature extraction and RMSD calculation relative to the native (input) ligand pose. The model is then trained to learn the relationship between pose-specific features and the corresponding RMSD of each docked pose. After training, we perform feature correlation analysis, model selection, and hyperparameter tuning. The final trained model predicts RMSD values for unseen ligand poses and re-ranks them based on lowest predicted RMSD, generating the final IRIS pose ranking.

The labeled features were then used to train IRIS to predict RMSD values for each pose and then re-rank the poses by their predicted RMSD. Finally, the trained model was applied to predict the RMSD for unseen poses, allowing for re-ranking of ligand poses based on their predicted RMSD, producing the final ranking.

The mean, median, and standard deviation of RMSD values for each rDock scoring function and search method combination were calculated for all NA-ligand complexes. The standard deviation (𝜎) is defined here as:

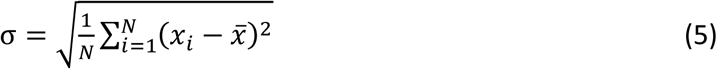

where 𝑁 is the total number of RMSD values, 𝑥_𝑖_ is the RMSD value for ligand pose 𝑖, and *x̄_i_* is the mean RMSD across ligand poses.

### Feature Importance

To determine the contributions of individual features to the predictions of the IRIS models, we performed a SHAP (SHapley Additive exPlanations) analysis^54^. SHAP values quantify the contribution of each feature by distributing the difference between the actual prediction and the average prediction across all input features. The SHAP value (𝜙_𝑖_) for a given feature 𝑖 is defined as:

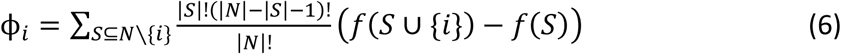

where *S* is a subset of all features excluding 𝑖, *f (S)* is the model output when using the feature subset *S*, 𝑁 is the total number of features, and *f (S ∪* {*i*}) is the model output when adding feature 𝑖 to subset *S*. The term 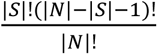 represents the weighting factor used to fairly distribute feature contributions across all possible subsets.

*Distribution of Errors.* We calculated the distribution of errors between predicted and actual RMSD values across the test set. The bias, *B*, of the model is defined as:

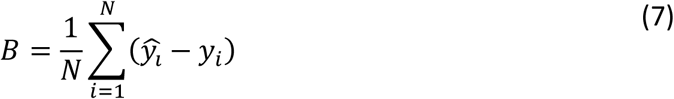

where 𝑁 is the number of ligand poses, *ŷ_i_* is the predicted RMSD for a given pose, and 𝑦_𝑖_ is the actual RMSD for the same pose. Bias quantifies the average difference between predicted and actual values, indicating whether the model systematically overestimates or underestimates RMSD across test set complexes. A bias close to 0 suggests minimal systematic error. The variance (*σ*^2^) of the prediction errors is given by:

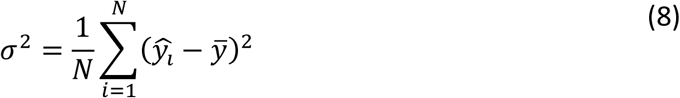

where *ȳ* is the mean predicted RMSD of the complex. Variance measures the spread of individual prediction errors, with higher values indicating greater inconsistency in the ability of the model to predict RMSDs for different complexes.

### RNAPosers comparison

We evaluated the performance of RNAPosers in reproducing near-native poses for all the RNA-small molecule complexes used in this work. The publicly available code for RNAPosers in the master version at https://github.com/atfrank/RNAPosers was used. Receptor Mol2 files, ligand SDF files, and the rDock output file containing ligand poses were given as inputs to the program.

For each ligand atom in the corresponding SDF file, RNAPosers generates a vector (or fingerprint) by calculating the distances of the ligand atom to all RNA atoms that are within 20 Å. These distances are then grouped by ligand–RNA atom pair types. The final fingerprint for each pose reflects the overall spatial pattern of ligand–RNA interactions^29^. RNAPosers also includes scoring terms from rDock, with the final model using both the fingerprints and rDock terms as input features^29^.

## Results

We initially assessed the performance of the rDock scoring functions *dock* and *dock_solv* (see Methods) on the constructed dataset of 1,435 nucleic acid-ligand (NA-ligand) complexes (**Table 1)**. Both scoring functions were tested using both the *RL* and *TS* docking methods.

**Table 1.** Comparison of the rDock default scoring functions (*dock* and *dock_solv*) and search methods (Two Sphere (*TS*) and Reference Ligand (*RL*)) on pose prediction accuracy. All mean, median and standard deviation values are in Å.

**Top Ranked**
|  | Method | Mean | Median | Stdev | % Poses < 2.0 Å RMSD |
| --- | --- | --- | --- | --- | --- |
| 0 | *RL_dock* | 4.4 | 3.6 | 3.3 | 29.0 |
| 1 | *RL_dock_solv* | 4.3 | 3.3 | 3.4 | 32.1 |
| 2 | *TS_dock* | 7.2 | 7.3 | 4.1 | 15.0 |
| 3 | *TS_dock_solv* | 7.1 | 7.3 | 4.2 | 16.4 |

**Best of Top 5**
|  | Method | Mean | Median | Stdev | % Poses < 2.0 Å RMSD |
| --- | --- | --- | --- | --- | --- |
| 0 | *RL_dock* | 3.1 | 2.5 | 2.5 | 40.8 |
| 1 | *RL_dock_solv* | 2.9 | 2.3 | 2.5 | 44.6 |
| 2 | *TS_dock* | 5.9 | 5.9 | 3.9 | 22.0 |
| 3 | *TS_dock_solv* | 5.9 | 5.9 | 4.0 | 22.3 |

**Best of 100**
|  | Method | Mean | Median | Stdev | % Poses < 2.0 Å RMSD |
| --- | --- | --- | --- | --- | --- |
| 0 | *RL_dock* | 1.5 | 1.3 | 0.9 | 77.0 |
| 1 | *RL_dock_solv* | 1.5 | 1.2 | 1.0 | 76.4 |
| 2 | *TS_dock* | 4.1 | 3.1 | 3.4 | 37.2 |
| 3 | *TS_dock_solv* | 4.1 | 3.0 | 3.4 | 37.8 |

Performance was measured by first determining the mean, median, and standard deviation of RMSDs of three groups:

1. the top-ranked ligand pose per complex (“Top Ranked”),
2. the lowest RMSD pose among the top 5 ranked ligand poses per complex (“Best of Top 5”), and
3. the lowest RMSD ligand pose found among all 100 generated poses per complex (“Best of 100”).

Additionally, we report the percentage of poses achieving RMSD values below 2.0 Å for each group, which is a common threshold used in docking studies for acceptable prediction accuracy ^11,27^, often known as the success rate. RMSD distributions across these criteria are visually presented in **Figure S2**.

The results are consistent with previous analyses of rDock pose prediction accuracy ^11,27^. As expected, across all metrics the *RL* method consistently outperforms the *TS* method. For the *Top Ranked* pose, *RL_dock_solv* achieves the highest accuracy, with 32.1% of poses under 2.0 Å

RMSD. When considering the *Best of Top 5*, the RL methods again demonstrated superior performance, *RL_dock_solv*, for example, increasing the percentage of poses below 2.0 Å to 44.6%. For the *Best of 100* across all complexes, both *RL_dock* and *RL_dock_solv* identified near- native poses in ∼77% of cases, with *RL_dock* performing slightly better than *RL_dock_solv*.

Following the above comparative assessment of the rDock native scoring functions, we aimed to improve pose ranking accuracy using ML. All further analysis focuses on improving the pose ranking accuracy of the *RL_dock* method, as it provided the highest number of complexes of the 100 generated with at least one pose below 2.0 Å RMSD and thus presents the greatest potential for successful re-ranking.

For the machine learning analysis, the dataset was split into training and test sets as detailed in the Methods section. In 79.4% of the test set complexes (i.e., 112 out of 141), RL_dock was able to generate at least one pose among the 100 poses per complex with an RMSD below 2.0 Å. (This differs from the percentage reported for the full dataset in **Table 1**, which includes both training and test complexes.) The challenge for ML, then, is to develop a method that can effectively re-rank poses within these 112 complexes. All subsequent values are thus expressed as a percentage of these 112 complexes, 100% being the best possible success rate that would be in principle achievable, through perfect ML ranking, given the available poses are generated by rDock.

A comprehensive set of features was generated and calculated for each of the 100 poses per complex as described in Methods. These features served as inputs for regression models aimed at predicting the RMSD of each pose relative to the native pose. The goal was to rank ligand poses based on predicted RMSD, with lower predicted RMSD values indicating better poses. To systematically evaluate the performance of a wide variety of different regression algorithms for RNA–ligand pose re-ranking, we implemented a benchmarking framework using K-fold cross-validation applied to the training set. We assembled a model zoo consisting of 13 widely used regressors, selected to represent diverse modeling strategies including linear, ensemble, kernel-based, and tree-based methods. The models in the zoo included Ridge^55^, Lasso^56^, ElasticNet^57^, Bayesian Ridge^58^, Support Vector Regression (SVR)^59^, LinearSVR^60^, K-Nearest Neighbors (KNN)^61^, Decision Tree^62^, Random Forest^63^, Gradient Boosting^64^, AdaBoost^65^, Bagging^66^, and XGBoost^67^. Default sci-kit learn^68^ hyperparameters were used for all models in the model zoo.

To ensure robust performance estimates, we performed K-fold cross-validation using five folds on the training set only, leaving the held-out test set out of all splits to avoid information leakage. For each model, we recorded the mean absolute error (MAE) and mean squared error (MSE) across all five folds, along with their standard deviations. The model with the lowest MAE was designated as the best model in the model zoo. This uniform evaluation strategy enabled direct comparison across models and informed selection of the model to continue toward hyperparameter tuning. The MAE and MSE of the model zoo across all tested models are presented in Table S4.

Based on the results from the model zoo, the Random Forest Regressor was selected as the IRIS regression model. We then conducted hyperparameter tuning on the Random Forest model using K-fold cross-validation with 5-folds. A randomized search was used to optimize model hyperparameters across the hyperparameter space presented in Table S5, with the goal of minimizing the MAE across folds.

The best-performing configuration was selected based on cross-validated MAE, and the final IRIS model was retrained on the full training set using these parameters. For completeness, we also calculated the mean squared error (MSE) to characterize the distribution of residuals. These loss functions are defined as:

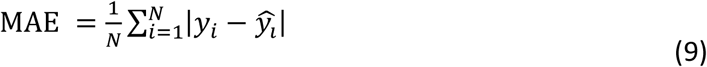

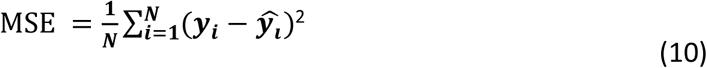

where 𝑁 is the total number of ligand poses, 𝑦_𝑖_ is the actual heavy atom RMSD of pose 𝑖 relative to the experimental structure, *ŷ_i_* is the predicted RMSD for pose *i*, and *ȳ_i_* is the mean of actual RMSD values. MAE quantifies the average absolute difference between predicted RMSD of each pose and actual RMSD of each pose across all 100 poses per complex, with lower values indicating better performance, while MSE provides a squared penalty for large deviations, making it more sensitive to outlier predictions.

Next, we compared the ability of the IRIS model to identify near-native poses (RMSD < 2.0 Å) on the *RL_dock* test set with that of the rDock native scoring function. Additionally, to determine whether a more complex model such as IRIS is necessary for improved pose ranking, we also evaluated whether the internal scoring terms computed by rDock alone are sufficient to predict pose quality. Specifically, we trained a linear regression^69^ model to predict RMSD using only the original rDock score components. This model serves as a reweighted variant of the rDock scoring function, providing an optimized linear combination of terms already available during docking. We further benchmarked IRIS against RNAPosers, a recently published RNA-specific pose re-ranking method. All four models—the default rDock scoring function, the reweighted rDock model, RNAPosers, and IRIS—were evaluated on the same *RL_dock* test set (**Figure 3**).

**Figure 3.**
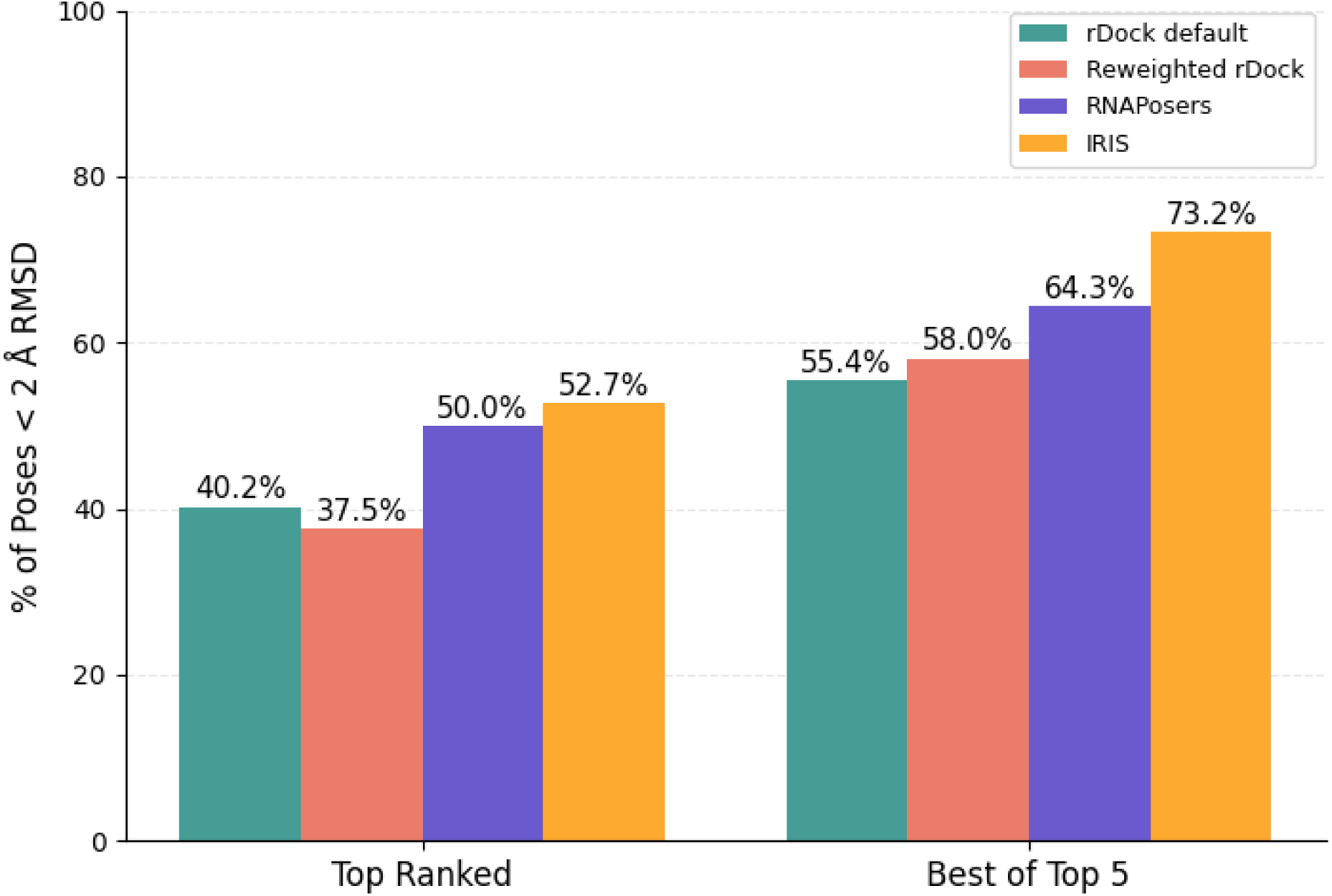
Normalized performance of IRIS re-ranking (yellow), RNAPosers (purple), reweighted rDock scoring function (red), and the standard rDock default scoring function (teal) on the RL_dock test set. Performance was evaluated as the percentage of test set complexes in which the Top Ranked pose (ranked #1 by each method) or the Best of Top 5 poses (lowest RMSD among top five ranked poses) achieved an RMSD ≤ 2.0 Å. All values are normalized to the maximum achievable success rate for RL_dock (79.4%), which reflects the percentage of test set complexes for which rDock generated at least one of the 100 generated poses within 2.0 Å RMSD of the experimental structure.

Using the rDock default scoring function, the Top Ranked pose was below 2.0 Å in 40.2% of cases. Reweighting the rDock scoring terms decreased this to 37.5%. RNAPosers achieved 50.0%, while re-ranking with IRIS increased this to 52.7%. For the Best of Top 5, rDock achieved 55.4%, the reweighted model reached 58.0%, RNAPosers achieved 64.3%, and IRIS achieved 73.2%. These results show that while the rDock scoring terms contain some predictive signal, a simple linear reweighting is not sufficient to substantially improve performance, and IRIS performs better than both native and reweighted rDock. Additionally, while RNAposers approached IRIS in Top-1 success, IRIS provided a much larger advantage in the Best of Top 5 setting. IRIS provides the greatest improvement in re-ranking accuracy and approaches the theoretical upper limit of performance defined by the quality of poses generated by rDock.

To complement the quantitative evaluation, we examined the structural features of complexes where IRIS predictions deviated most strongly from the true top-ranked pose. We defined deviation as the difference between the actual rank of the IRIS-predicted top-1 pose and the true top-1 pose. In several test cases, including 1FHY, 1M69_2, and 7WII, the IRIS-predicted top-1 pose corresponded to one of the lowest-ranked poses by RMSD (actual ranks 100, 99, and 98, respectively). Visual inspection of these complexes revealed consistent challenges (**Figure S3**). In 1FHY, the ligand 4′-hydroxymethyl-4,5′,8-trimethylpsoralen is a rigid, polyaromatic intercalator bound within a DNA duplex. In 1M69_2, the ligand spermine is a highly flexible polyamine that binds diffusely along DNA grooves. In 7WII, the riboswitch ligand 7,8- dihydroneopterin presents a planar heteroaromatic scaffold capable of multiple stacking orientations within the RNA binding pocket.

Although chemically distinct, these ligands share properties that complicate pose re-ranking: either conformational plasticity (spermine) or multiple plausible stacking geometries (psoralens and pterins). Moreover, their binding environments—extended DNA grooves or relatively shallow RNA pockets—provide fewer geometric constraints than compact RNA pockets. These conditions increase the number of near-degenerate poses, making it more difficult for IRIS to reliably identify the true near-native pose.

To assess how performance varied by nucleic acid type, we analyzed the RNA and DNA complexes within the test set separately. Of the 141 test set complexes, 108 were RNA and 33 were DNA. For each subset, we first determined the proportion of complexes for which rDock was able to generate at least one pose within 2.0 Å RMSD—representing the theoretical upper limit of re-ranking success. This maximum achievable success rate was 82.41% for RNA (89 of 108 complexes) and 69.70% for DNA (23 of 33 complexes). All subsequent model success rates were normalized to these respective ceilings to enable fair comparison.

IRIS consistently outperformed both baselines on the RNA subset (**Figure 4**). Using the rDock default scoring function, the normalized Top Ranked success rate was 43.8%. Reweighting the rDock scoring terms decreased this to 41.6%. Re-ranking with RNAposers improved this to 55.1%, while IRIS further increased it to 57.3%. For the normalized Best of Top 5 success rates, rDock achieved 59.5%, the reweighted model reached 61.8%, RNAposers achieved 69.7%, and IRIS achieved 78.6%. These results show that IRIS outperforms all tested methods for RNA pose re-ranking and, similar to the overall performance, while RNAposers rivals IRIS in Top-1 ranked accuracy, IRIS held a clear advantage in recovering near-native poses within the top five ranked candidates.

**Figure 4.**
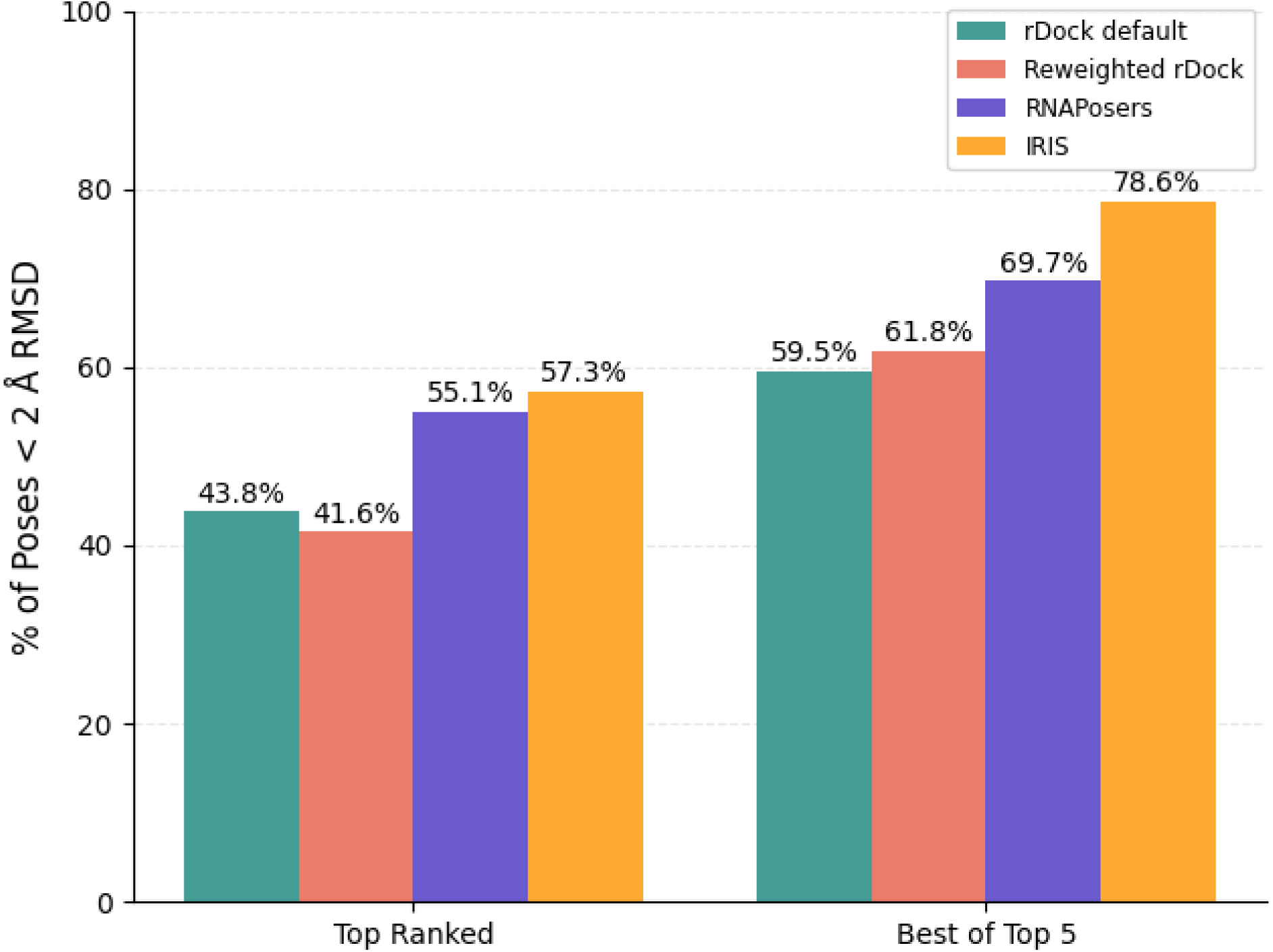
Normalized performance of IRIS re-ranking (gold), RNAPosers (purple), reweighted rDock scoring function (coral), and the standard rDock default scoring function (teal) on the RNA subset of the RL_dock test set. Performance was evaluated as the percentage of RNA test set complexes in which the Top Ranked pose (ranked #1 by each method) or the Best of Top 5 poses (lowest RMSD among top five ranked poses) achieved an RMSD ≤ 2.0 Å. All values are normalized to the maximum achievable success rate for RNA (82.41%), which reflects the percentage of RNA test set complexes for which rDock generated at least one of the 100 poses within 2.0 Å RMSD of the experimental structure.

Results for the DNA subset were also computed (**Figure 5**). Using the default rDock scoring function, the normalized Top-1 success rate was 26.1%. Reweighting the rDock scoring terms decreased this slightly to 22.5%. RNAposers improved this to 30.4%, and IRIS further increased it to 34.8%. For the normalized Best of Top 5 success rates, rDock achieved 39.1%, the reweighted model reached 43.5%, RNAposers achieved 43.5%, and IRIS achieved 52.2%. These findings indicate that IRIS provides meaningful gains in pose selection accuracy across both RNA and DNA complexes, even when normalized to their respective upper bounds of docking success. As in the RNA subset, RNAposers approached IRIS in Top-1 success but failed to match IRIS’s larger gains in the Best of Top 5 performance.

**Figure 5.**
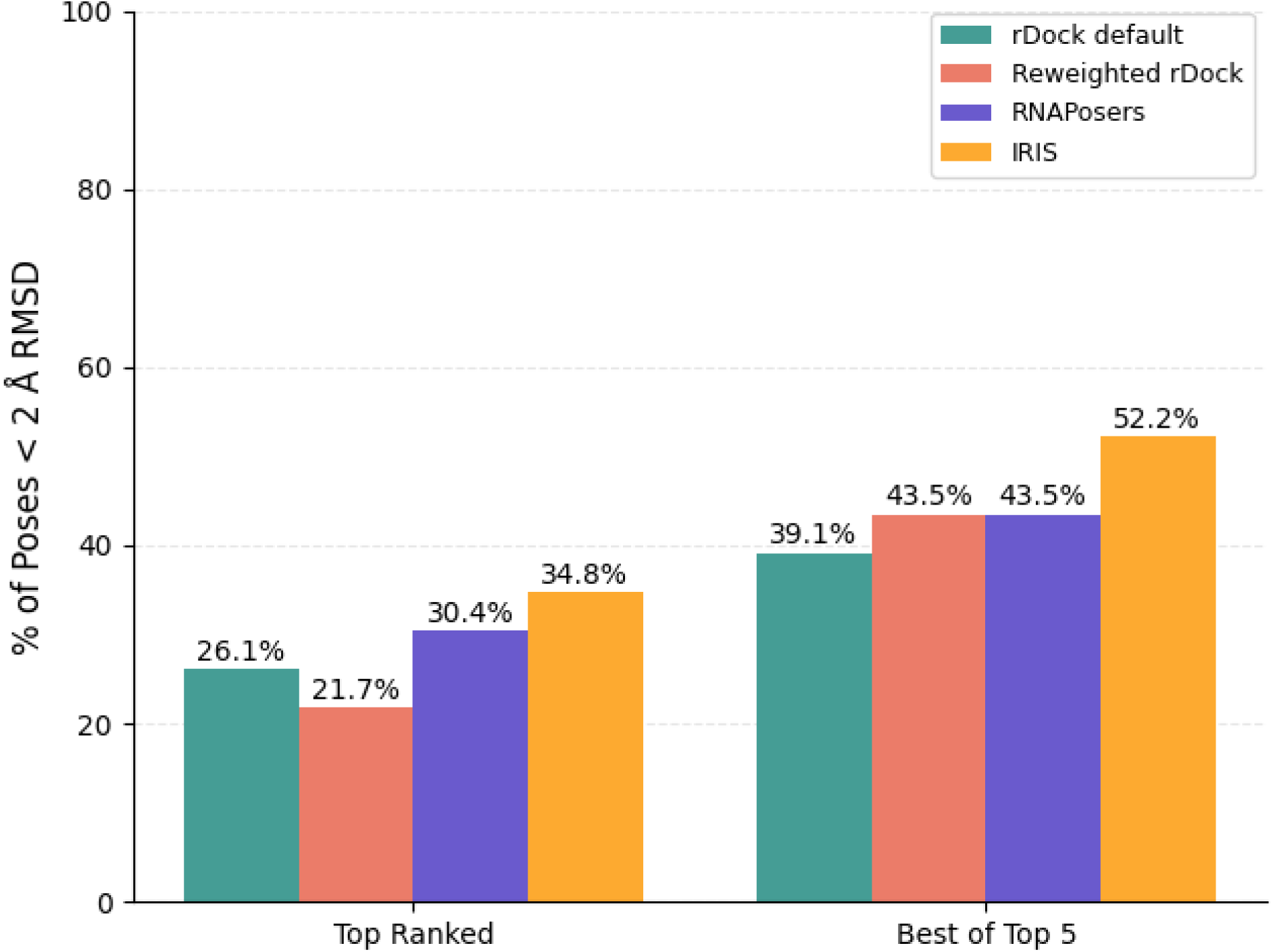
Normalized performance of IRIS re-ranking (gold), RNAPosers (purple), reweighted rDock scoring function (coral), and the standard rDock default scoring function (teal) on the DNA subset of the RL_dock test set. Performance was evaluated as the percentage of DNA test set complexes in which the Top Ranked pose (ranked #1 by each method) or the Best of Top 5 poses (lowest RMSD among top five ranked poses) achieved an RMSD ≤ 2.0 Å. All values are normalized to the maximum achievable success rate for DNA (69.70%), which reflects the percentage of DNA test set complexes for which rDock generated at least one of the 100 poses within 2.0 Å RMSD of the experimental structure.

To address whether training on DNA- and RNA-ligand complexes provide some additional information which may influence model performance compared to training on solely RNA-ligand complexes, we retrained IRIS using only RNA complexes from the training set and evaluated it on the RNA subset of the test set. This was compared to the RNA subset performance of the IRIS model trained on all nucleic acid complexes (RNA+DNA) (**Figure S4**). For the RNA+DNA- trained model, the normalized Top Ranked and Best of Top 5 success rates on the RNA subset were 57.3% and 78.65%, respectively. Restricting both training and testing to RNA complexes reduced performance to 43.82% for Top Ranked and 75.28% for Best of Top 5. The decrease in performance across both metrics indicates that the IRIS model benefits from the inclusion of DNA complexes in training, likely because these structures expand the range of nucleic acid binding site geometries and interaction patterns encountered during learning. This broader structural and chemical diversity appears to help IRIS capture generalizable features of nucleic acid–ligand recognition, improving its ability to identify near-native poses even in RNA-specific evaluations.

**N**ext, to determine the contributions of individual features to the predictions of the IRIS models, we performed a SHAP (SHapley Additive exPlanations) analysis^54^. SHAP values quantify the contribution of each feature by distributing the difference between the actual prediction and the average prediction across all input features. Higher absolute SHAP values indicate features that strongly influence RMSD predictions. The mean SHAP value indicates the average directional impact a feature has across all predictions, showing whether a feature tends to increase or decrease the predicted RMSD. In contrast, the mean absolute SHAP value measures the average magnitude of contribution for a feature, regardless of direction, making it a reliable indicator of overall feature importance.

The top 10 features in the SHAP analysis, ranked by their mean absolute SHAP values, are shown in **Figure 6**, reflecting their overall influence on model performance. Notably, several rDock scoring terms, including SCORE.RESTR (0.2655), SCORE.INTER.norm (0.2199), and SCORE.norm (0.1357), rank among the most influential. While these terms alone fail to accurately rank poses (**Figure 3, 4, & 5**), their inclusion alongside diverse chemical and spatial descriptors enables the IRIS model to make more accurate and interpretable RMSD predictions.

**Figure 6.**
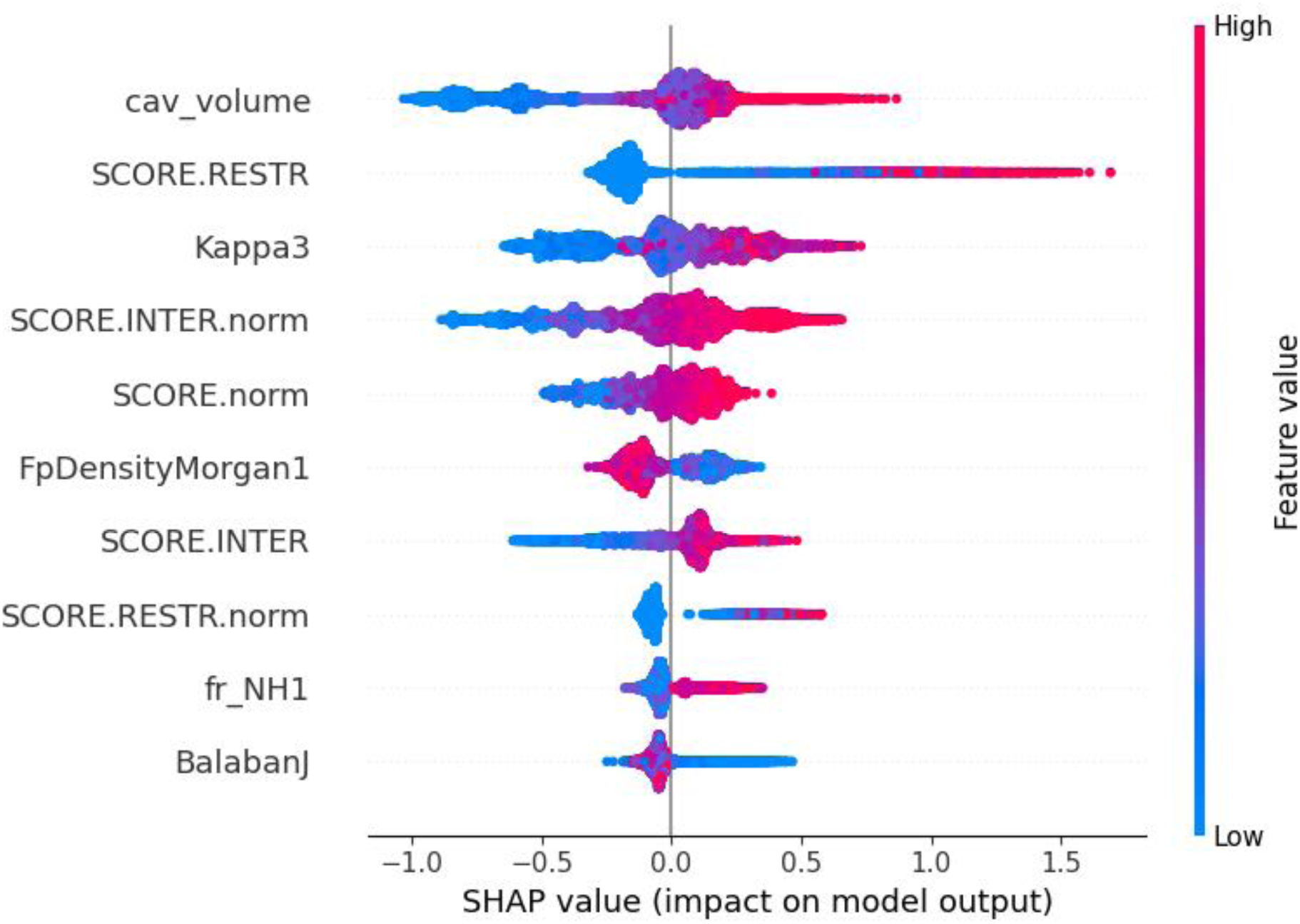
Top 10 most important features for the *RL_dock* IRIS model RMSD predictions: Each dot represents the corresponding feature describing a single ligand pose in the test set. The y-axis represents the original feature value for each pose relative to the values for the same feature, with colors describing whether the feature value is comparatively high (red) or comparatively low (blue).

Grouped together, the rDock-derived features in the SHAP analysis delineate distinct energetic contributions relevant to docking. SCORE.RESTR represents the penalty term for violating the spatial constraints of the defined docking cavity. In rDock, the cavity is specified by a set of grid points defining the permissible binding volume, and SCORE.RESTR increases when any ligand atom lies outside this volume, with the penalty scaled to the distance of the violation. The normalized version (SCORE.RESTR.norm) divides this penalty by the ligand’s heavy atom count to account for size effects^12,51^. SCORE.INTER and SCORE.INTER.norm quantify the ligand–receptor interaction energy, comprising van der Waals, polar (attractive and repulsive), and entropy-related terms, with the normalized form being divided by the ligand’s heavy atom count^12,51^. The SCORE.norm score is a heavy-atom-normalized version of the total docking score, combining all components (interaction, intramolecular strain, site-internal, and restraint energy) on a per-atom basis ^12,51^. In the IRIS model, higher SCORE.RESTR values correlate with increased predicted RMSD (mean SHAP = –0.0541), while more favorable interaction energies— SCORE.INTER.norm (mean SHAP = –0.0383), SCORE.INTER (-mean SHAP approx. –0.0415), and SCORE.norm (mean SHAP approx. –0.1029)—predicate lower RMSDs. Conversely, SCORE.RESTR.norm yields low mean SHAP impact (∼–0.0035), indicative of varied influence across docking scenarios. These patterns underscore the model’s capacity to interpret and contextualize energetic terms effectively, improving pose ranking beyond what any single raw or normalized score could achieve on its own.

Among the top 10 features, the cavity volume (cav_volume) emerged as the single most important feature (mean absolute SHAP = 0.2820, mean SHAP = –0.1427). This value is derived by calculating the convex hull volume of receptor atoms within 4 Å radius of the ligand, capturing the spatial openness of the binding pocket^51^. Larger cavities often correlate with higher predicted RMSD, likely due to increased conformational space the ligand can occupy. Kappa3, a molecular shape index that reflects the degree of linearity or compactness^70^, had a mean absolute SHAP of 0.2263. Higher values (more rod-like molecules) tended to reduce predicted RMSD (mean SHAP = –0.0214), suggesting that specific molecular shapes are more consistently placed during docking.

FpDensityMorgan1 (mean absolute SHAP = 0.1354) measures the local density of circular substructures within a Morgan fingerprint of radius 1, capturing fine-grained atomic environment complexity^71^. Higher values correlated with lower RMSD predictions, indicating that more densely featured local environments aid in pose recognition. BalabanJ, a topological descriptor reflecting global molecular connectivity^72^, had lower overall importance (mean absolute SHAP = 0.0709), but still trended negatively with RMSD (mean SHAP = –0.0293), suggesting modest stabilizing effects of optimal connectivity. fr_NH1 (mean absolute SHAP = 0.0731) reflects the number of primary amine groups capable of forming hydrogen bonds^35^, consistent with known preferences in RNA-ligand recognition. Its low mean SHAP value (–0.0054) suggests variable impact across complexes.

A complete list of feature definitions, sources, and calculation methods is provided in the Supporting Information, including detailed descriptions of all included energetic, geometric, and chemical descriptors.

Features in the SHAP analysis align closely with known determinants of RNA–ligand recognition. The Kappa3 shape index reflects the rod-like geometry of RNA-targeted molecules, a trait associated with enhanced stacking and groove fit^1^. FpDensityMorgan1, which measures local atomic environment complexity, captures the higher heteroatom content and functional group density common in RNA-binding ligands^73^. The fr_NH1 descriptor highlights the contribution of primary amines, which are frequently enriched in RNA binders due to their capacity to form hydrogen bonds and electrostatic contacts^74^. The inclusion of BalabanJ in the top 10 most important features supports the idea that increased molecular may be an important property of binding modes of ligands which target nucleic acids. Finally, cavity volume reflects the spatial constraints imposed by the binding pocket, where tighter volumes may limit ligand flexibility and encourage more consistent docking poses. Taken together, these shape- and complexity-related descriptors support the notion that RNA-binding ligands, which are often rod-like and rich in heteroatoms, achieve better fit within smaller, more confined cavities that constrain their degrees of freedom and enhance pose consistency^1,7383–86^

To assess the aggregate influence of related descriptors, features from the SHAP analysis were grouped into broader categories based on their origin and functional relevance: RDKit molecular descriptors, rDock sub-scores, ligand–receptor distances and geometries, binding site interactions, and RDKit fragment counts. The summed mean absolute SHAP values for each category were normalized to the total SHAP magnitude across all features to yield their relative contributions (**Figure S5**). RDKit molecular descriptors and rDock sub-scores contributed the largest shares (34.97 % and 32.64 %, respectively), highlighting the complementary roles of general ligand physicochemical properties and docking-derived energetic terms in driving RMSD predictions. Ligand–receptor distance and geometry metrics accounted for 20.72 % of total SHAP impact, indicating that spatial fit descriptors substantially influence pose ranking. Binding site interaction features contributed 7.68 %, while fragment count descriptors explained 4.00 %, reflecting their more limited influence. This category-level view shows that IRIS integrates multiple descriptor classes which supports the importance of combining docking scores with chemically and spatially diverse features. A more detailed description of which features are included in each category is given in the Supplementary Information.

Next, we assessed the sensitivity of the IRIS model to training data availability by constructing a learning curve using the IRIS training set (**Figure S6**). To ensure fair evaluation at each training size, we applied K-fold cross-validation with 5 folds. For each fold, the model was trained on a subsample of the training data and evaluated on the held-out fold, and this was repeated across ten incremental training set sizes. In this setup, 20% of the data is held out in each fold for validation, so even at the largest plotted training size the model is trained on only 80% of the available training data. This means the training MAE at the rightmost point of the curve is based on a 20% smaller dataset than the full training set used in fitting of the true IRIS model, which may contribute to the observed gap between training and validation errors. The shaded regions represent the standard deviation across folds and highlight variability due to data partitioning.

The resulting learning curve shows that training error increases modestly with more data. Meanwhile, validation error decreases steadily with increasing training size, indicating improving generalization. This trend contrasts with classic overfitting, in which validation error plateaus or increases as the model begins to memorize noise specific to the training data^75^. Even though the validation curve continues to decrease, since the training and validation curves do not fully converge, it suggests the IRIS model remains data-limited, and performance could likely be enhanced with additional structurally diverse NA-ligand complexes in the training set.

To assess the consistency of IRIS predictions across ligand poses, we calculated the mean absolute error (MAE) between predicted and actual RMSD values for each complex in the test set (**Figure 7**). Each complex includes approximately 100 poses, allowing us to evaluate how prediction errors vary across the ligand ensemble. Complexes with high MAE exhibit larger deviations between predicted and actual RMSD values across poses, reflecting inconsistent or poor ranking performance. For example, complex 453d (complex 1 in **Figure 7**) displays the highest average prediction error at 5.35 Å, with a standard deviation of 0.44 Å, indicating high error magnitude despite relatively low variability across poses. In contrast, complex 1m69_2 (complex 141 in **Figure 7**) shows the lowest MAE at 0.35 Å with a standard deviation of 0.34 Å, suggesting consistently accurate predictions across all poses.

**Figure 7.**
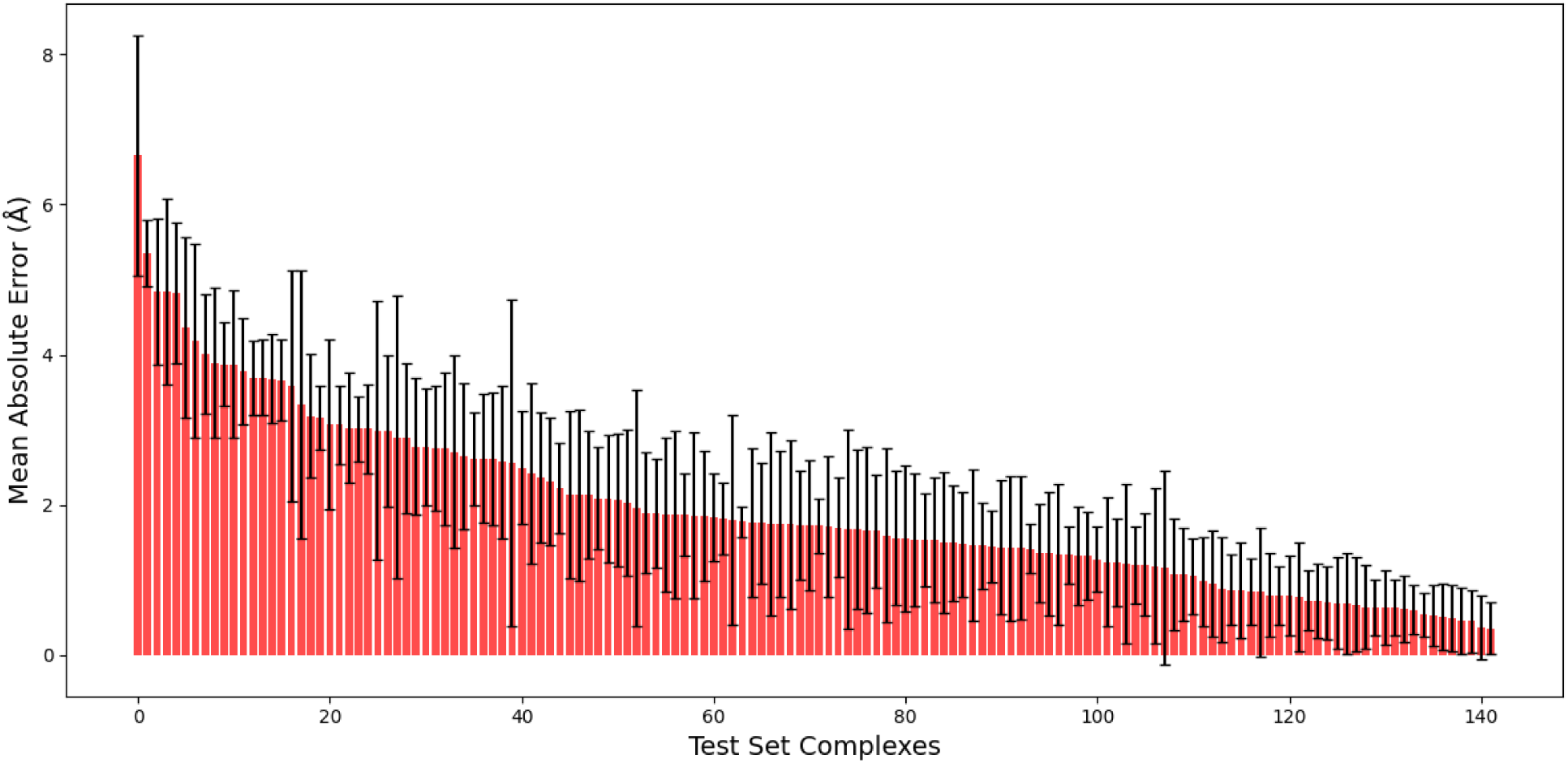
Mean Absolute Error (Å) in prediction accuracy across test set complexes. The x-axis represents test set complexes sorted in descending order of MAE, while the y-axis shows the MAE of predicted RMSD values for all ligand poses within each complex. Error bars represent one standard deviation above and below the mean MAE, reflecting the variability in prediction accuracy across poses within each complex.

To better understand the molecular factors associated with prediction error, we compared ligand features between the top 10 complexes with the highest and lowest per-complex MAE (**Table 2**). For each group, the mean and standard deviation of molecular weight, number of rotatable bonds, and cavity volume are reported.

**Table 2.** Comparison of average ligand features for test complexes with the top 10 highest vs. top 10 lowest mean absolute error (Å) values. For each group, the mean and standard deviation of each ligand feature are reported.

| Descriptor | High MAE (Mean) | High MAE (Stdev) | Low MAE (Mean) | Low MAE (Stdev) |
| --- | --- | --- | --- | --- |
| Molecular Weight (Da) | 527.7 | 171.0 | 127.2 | 75.2 |
| Number of Rotatable Bonds | 8.0 | 4.1 | 1.8 | 1.5 |
| Cavity Volume (Å<sup>3</sup>) | 1294.5 | 436.2 | 428.4 | 185.9 |

Complexes with the highest MAE values tended to contain substantially larger and more flexible ligands than those with the lowest MAE values, as reflected by higher mean molecular weight (527.7 Da vs. 127.2 Da) and greater numbers of rotatable bonds (8.0 vs. 1.8). These ligands were also bound in substantially larger cavities on average (1294.5 Å³ vs. 428.4 Å³), providing greater conformational space and potentially increasing the difficulty of accurately predicting pose RMSD. In contrast, low-MAE complexes generally involved smaller, more rigid ligands in more spatially constrained binding sites, which may facilitate more consistent pose prediction.

We then sought to visually analyze the top three complexes in the highest and lowest MAE groups. To this end, we visualize each of these complexes using PyMOL^36^ to identify similarities or differences that may contribute to their respective error ranges in pose RMSD prediction (**Figure S7 & S8**). For clarity, each visualization retains only the nucleic acid receptor (RNA or DNA) bound to the ligand and the ligand itself, with all other structural components removed to focus on the nucleic-acid binding pocket.

It is important to note that the complexes highlighted here do not directly reflect the ability of IRIS to identify the correct top-ranked pose, but rather its ability to predict RMSD values consistently across all poses for a given complex. A complex may exhibit a high per-complex MAE because its ensemble of poses spans a wide structural range—even if the top-ranked pose is close to the native structure—while a low per-complex MAE indicates that predicted RMSDs are consistently accurate across poses with minimal variability. This distinction is essential when interpreting MAE-based comparisons, as they evaluate prediction consistency rather than pose-ranking success alone.

The three highest-MAE complexes all contain large, heteroaromatic, drug-like ligands in complex with DNA. These ligands have elongated or curvilinear architectures, multiple aromatic rings, and high numbers of rotatable bonds. 1K2Z^76^ (MAE = 4.83 Å ± 1.24 Å) features distamycin A, a polyamide antibiotic that binds in the minor groove of AT-rich DNA and contains multiple hydrogen bond donors/acceptors. 3EUI^77^ (MAE = 4.84 Å ± 0.97 Å) contains a bulky 3,6- disubstituted acridine derivative (543 Da) with two cationic 2-ethylpiperidine side chains that π- stack on the terminal G-quartet of a telomeric DNA G-quadruplex. 453D^78^ (MAE = 5.35 Å ± 0.44 Å) holds a symmetric bis-benzimidazole analogue that binds in the DNA minor groove at AATT sites. For each complex, we include all 100 poses generated by rDock in the MAE calculation. In ligands with many degrees of freedom, conformational flexibility permits a wide range of energetically plausible binding poses, producing greater structural dispersion at and around the binding site. This increased pose variability translates into larger RMSD differences across the ensemble, elevating both the mean and standard deviation of prediction errors even if the top pose is below 2 Å RMSD to the native pose.

The three lowest-MAE complexes contain small, low-complexity ligands with compact shapes, short molecular lengths, and minimal rotatable bonds. 2IBK^79^ (MAE = 0.61 Å ± 0.45 Å) is bound here to ethylene glycol. Ethylene glycol is commonly used in protein crystallography as a cryoprotectant to prevent ice formation and preserve crystal integrity during flash-cooling^80^. 5BJP^81^ (MAE = 0.54 Å ± 0.29 Å) contains dimethyl sulfoxide as the small molecule depicted here. Dimethyl sulfoxide is often used as a cryoprotectant during X-ray crystallography^82^. 6D8A^83^ (MAE = 0.61 Å ± 0.33 Å) has (4S)-2-methylpentane-2,4-diol, a small chiral glycol used as a crystallization additive^84^, which is bound to a DNA strand in from *Rhodobacter sphaeroides*. As with the high-MAE group, these MAE values are based on all 100 rDock-generated poses per complex. However, the minimal conformational freedom of these ligands constrains the set of plausible binding orientations, reducing variability at the binding site and leading to consistently low RMSD prediction errors across the ensemble.

## Conclusions

Building upon previous ML-based methods^28–30^ for improving pose accuracy in RNA- ligand docking, we present here IRIS, an ML-based framework designed to improve RNA-ligand docking pose ranking by reranking rDock-generated poses. In the test set constructed, IRIS consistently ranks a greater proportion of near-native poses below 2.0 Å RMSD than the standard rDock scoring function, the linear regression model which uses the rDock scoring function terms, and the RNAPosers scoring function. SHAP analysis shows that both rDock scoring function terms and ligand-specific physicochemical properties are important to pose ranking, suggesting that future ML-based scoring approaches should focus on selection and weighting of these descriptors.

The downward trend in the learning curve (**Figure S6**indicates that the scale of training data influences the effectiveness of the ML-based re-scoring method. Therefore, the inclusion of a larger and more diverse set of nucleic acid-ligand complexes likely contributed to the improved ranking accuracy in IRIS compared to previous methods. This finding is consistent with previous studies, which have demonstrated that the composition of the training dataset influences the performance of ML models used for molecular property prediction^85,86^.

By identifying poses that are structurally closer to the experimentally relevant binding mode, IRIS may be useful in increasing the probability that subsequent affinity calculations— whether through scoring functions, molecular dynamics simulations, or free energy perturbation methods—are based on physically meaningful conformations rather than misaligned or non-physiological poses. Our pose re-ranking tool is particularly valuable in virtual high-throughput screening (VHTS), where docking algorithms generate a set of poses for each ligand, but their scoring functions do not always rank the near-native pose at the binding site correctly^87^. Without effective pose filtering, inaccurate poses can negatively impact the accuracy of affinity predictions, as illustrated by the influence of the ligand binding pose on protein affinity predictions^88^. By prioritizing near-native poses, IRIS helps focus computational efforts towards ligands with a higher probability of genuine binding, thereby improving the overall efficiency and accuracy of lead selection in RNA-targeted drug discovery.

The limited availability of large datasets remains a challenge for many ML methods aimed at predicting chemical properties^89^. IRIS accuracy is limited by the size of the training dataset, and additional experimentally determined RNA-ligand complexes are likely to improve generalization. Model accuracy is also found to decrease for ligands with higher molecular weight, more rotatable bonds, and larger cavity volume, indicating challenges in ranking highly flexible and bulky ligands. This is consistent with previous studies which observed that larger ligands with more rotatable bonds tend to correspond with greater inconsistency and decreased accuracy in protein-ligand docking pose predictions^90^. Other limitations of IRIS are that it is strictly a re-ranking tool, meaning its performance depends on the generation of near-native poses by rDock, and it does not perform pose refinement.

Considerations for future work include extending IRIS beyond pose re-ranking to predict binding affinities. This might be achieved by leveraging current IRIS features but integrating experimental binding data and training the model on affinity metrics, such as dissociation constants, for complexes for which such values have been experimentally determined. Moreover, expanding IRIS to function independently of rDock search constraints would enhance its ease of use, allowing it to generalize across diverse docking pipelines. Adapting IRIS for use with other docking programs is straightforward and would further broaden its applicability.

## Supporting information

transit_high_tanimoto_clusters

IRIS_redundant_pairs

shap_feature_to_category

Feature_set

iris_train_test_split

NA_complexes

## Acknowledgement

We are grateful to Dr. Anna Pyle for helpful discussions.

## Author contributions

Andrew Amburn (Conceptualization, Methodology, Formal analysis, Software, Methodology, Visualization, Writing—original draft, Data curation), Shalini J. Rukmani (Formal analysis, Data curation, Resources, Writing—review & editing), Jerry M. Parks (Conceptualization, Supervision, Resources, Writing—review & editing), and Jeremy C. Smith (Conceptualization, Supervision, Resources, Writing—review & editing)

## Supplementary data

Supplementary data is available at NAR online.

## Data and Software Availability

The IRIS software is freely available at https://github.com/AndrewAmburn/IRIS. The data that support the findings of this study are available at https://zenodo.org/records/15597920.

