## Supplementary_information for "IRIS: A Machine Learning-Based Pose Re-Ranking Tool for RNA-Ligand Docking"

[a]The Bredesen Center for Interdisciplinary Research and Graduate Education, University of Tennessee, Knoxville, Tennessee, [b]UT/ORNL Center for Molecular Biophysics, Oak Ridge National Laboratory, Oak Ridge, TN, United States, [c]Department of Biochemistry and Cellular and Molecular Biology, University of Tennessee, Knoxville, TN, United States, [d]Biosciences Division, Oak Ridge National Laboratory, Oak Ridge, TN, USA,

**Table S1**

**Table S1**. Ligand clusters with the highest frequency in the training set. Clusters here are defined as 2 or more ligands with greater than 0.8 Tanimoto similarity.

| Cluster ID | % of Training Set |
|---|---|
| 12 | 4.07% |
| 64 | 2.04% |
| 45 | 1.80% |
| 8 | 1.56% |
| 23 | 1.20% |
| 13 | 1.20% |
| 10 | 1.08% |
| 16 | 0.96% |
| 7 | 0.96% |
| 84 | 0.84% |

### Table S2

**Table S2**. Test complexes with maximum common substructure (MCS) RMSD ≤ 5 Å to a training complex. Whether the test complex was DNA or RNA is included.

| Test Complex | Train Complex | MCS RMSD (Å) | NA Type |
|---|---|---|---|
| 1xvn | 1xvk | 0.35 | DNA |
| 1ekh | 1eki | 3.59 | DNA |
| 3m8r_3 | 3m8s | 1.00 | DNA |
| 2ktz | 2ku0 | 3.58 | RNA |
| 6tf1 | 6tb7 | 0.70 | RNA |

### Table S3

**Table S3**. Test set ligand pairs with Tanimoto similarity ≥ 0.8 and their MCS RMSD values. Among the 29 high-similarity ligand pairs identified in the test set (Tanimoto coefficient ≥ 0.8), we computed the root-mean-square deviation (RMSD) between their maximum common substructures (MCS) using RDKit, without alignment or minimization. Despite strong 2D similarity (including many Tanimoto values of 1.0), all pairs exhibited large MCS RMSD values—ranging from 8 Å to over 125 Å—indicating substantial spatial diversity. This analysis confirms that the test ligands are not only chemically diverse but also structurally distinct in 3D where chemically similar, supporting the validity of the test set for evaluating pose re-ranking performance.

| **Ligand1** | **Ligand2** | **Tanimoto coefficient** | **MCS RMSD** |
|---|---|---|---|
| 1dcr | 2o3y | 1 | 8.467 |
| 258d_2 | 1dcr | 1 | 14.921 |
| 258d_2 | 2o3y | 1 | 17.837 |
| 2o3y | 2tra | 1 | 19.204 |
| 2a5r | 6jjh | 1 | 19.61 |
| 1i2y | 1vro | 1 | 22.012 |
| 1vro | 1dcr | 1 | 24.414 |
| 1dcr | 2tra | 1 | 24.416 |
| 258d_2 | 1i2y | 1 | 24.42 |
| 1i2y | 1dcr | 1 | 26.09 |
| 258d_2 | 2tra | 1 | 28.247 |
| 1i2y | 2o3y | 1 | 29.693 |
| 1vro | 2o3y | 1 | 30.176 |
| 258d_2 | 1vro | 1 | 30.255 |
| 7lng | 6u7z | 1 | 30.828 |
| 1i2y | 2tra | 1 | 31.096 |
| 6hbx | 5o62 | 1 | 34.185 |
| 1dns | 1m69_2 | 1 | 40.474 |
| 1vro | 2tra | 1 | 40.835 |
| 5z71 | 1mwl | 1 | 42.479 |
| 6d8a | 1i7j | 1 | 48.581 |
| 4kqy | 2qwy | 1 | 58.471 |
| 6hag | 7dwh | 0.8028 | 67.853 |
| 4frg | 4frn | 1 | 91.736 |
| 5d5l | 2miy | 1 | 105.697 |
| 6hag | 3npn | 0.8551 | 125.121 |
| 8swo | 8sx5 | 1 | 126.84 |
| 3npn | 7dwh | 0.9091 | 127.675 |

**Figure S1**

**Figure S1.** All pairwise Tanimoto coefficients between the 141 test ligands using Morgan fingerprints (radius = 2, 2048 bits) were computed to assess ligand chemical diversity. The resulting similarity matrix revealed that most ligand pairs had low structural similarity. Only 29 ligand pairs exhibited a Tanimoto coefficient ≥ 0.8, confirming that the test set covers a broad and chemically diverse range of 2D fingerprint space.

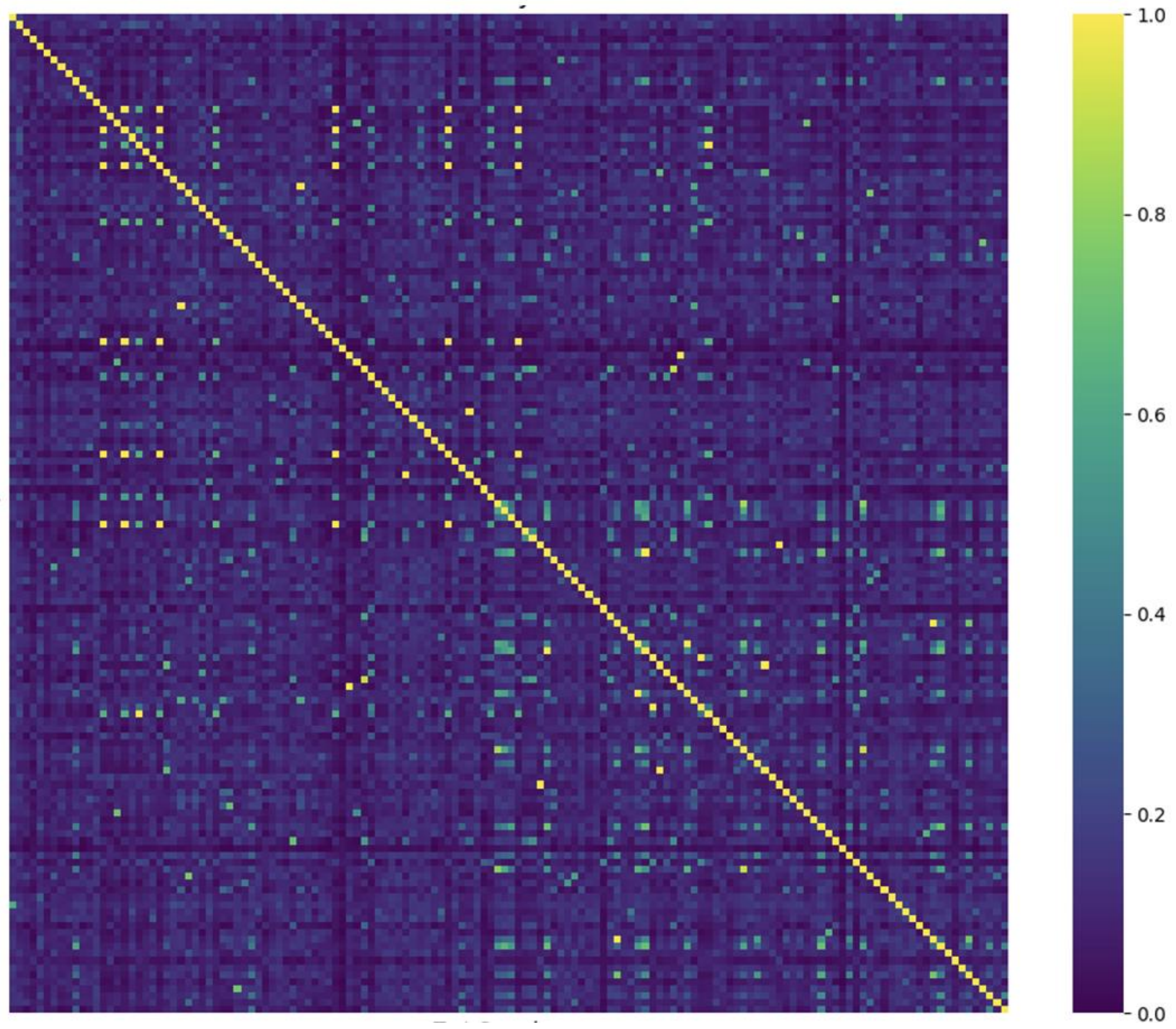

**Figure S2.**

**Figure S2**. Comparison of RMSD distributions in the full dataset, including training and testing sets, obtained with the dock and dock_solv scoring functions for (**A**) the top-ranked ligand, the (**B**) lowest RMSD pose out of the top five ranked ligands, and (**C**) the lowest RMSD pose out of 100 poses.

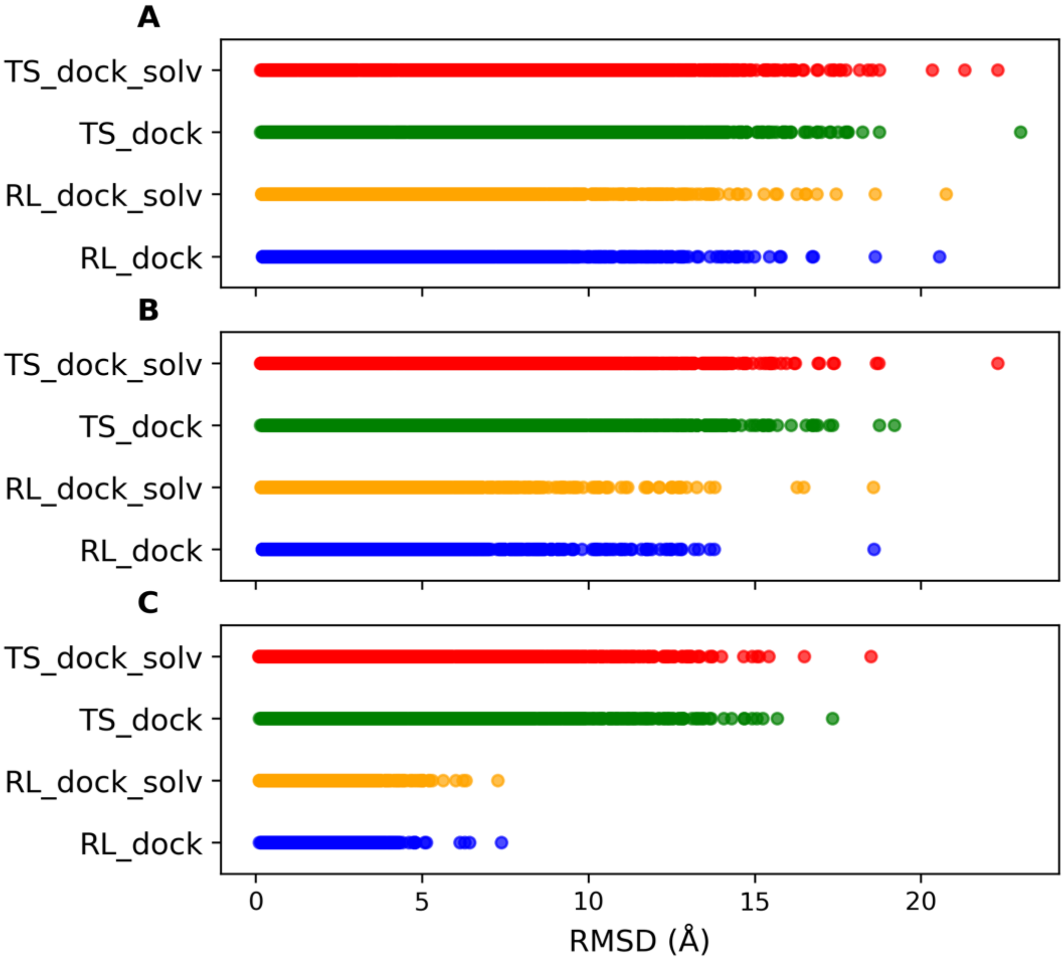

**Table S4.**

**Table S4.** Mean absolute error (Å) and mean squared error ($Å^2$) values for all tested models in the model zoo for IRIS**.**

| Model | MAE (Å) | MAE Std (Å) | MSE (Å²) | MSE Std (Å²) |
|---|---|---|---|---|
| **RandomForest** | **1.342** | **0.012** | **4.390** | **0.083** |
| Bagging | 1.374 | 0.011 | 4.888 | 0.072 |
| DecisionTree | 1.506 | 0.004 | 8.892 | 0.178 |
| KNN | 1.515 | 0.014 | 5.962 | 0.108 |
| XGBoost | 1.692 | 0.011 | 5.473 | 0.088 |
| SVR | 1.911 | 0.015 | 8.919 | 0.168 |
| GradientBoosting | 2.213 | 0.010 | 7.566 | 0.116 |
| LinearSVR | 2.241 | 0.011 | 11.676 | 0.954 |
| Ridge | 2.373 | 0.010 | 8.654 | 0.090 |
| BayesianRidge | 2.374 | 0.010 | 8.650 | 0.078 |
| ElasticNet | 2.684 | 0.011 | 11.061 | 0.144 |
| AdaBoost | 2.705 | 0.009 | 10.094 | 0.128 |
| Lasso | 2.800 | 0.012 | 12.178 | 0.168 |

**Table S5.**

**Table S5.** Hyperparameter search space for the Random Forest regressor used in IRIS. RandomizedSearchCV was used with grouped 5-fold cross-validation and mean absolute error as the scoring metric.

| Hyperparameter | Search Space | Notes |
|---|---|---|
| max_depth | 5, 10, 20, 30, 40, None | Maximum depth of trees |
| min_samples_leaf | 1, 3, 5 | Minimum samples per leaf node |
| max_features | “sqrt”, “log2”, 0.5, None | Features considered per split |
| n_estimators | 50, 100, 200, 500, 1000 | Number of trees in the forest |

**Figure S3**

**Figure S3.** Structural visualization of the three worst-predicted test set complexes by IRIS. Each panel shows the nucleic acid receptor (white cartoon) bound to the IRIS-predicted top-1 ligand pose (colored sticks). For these complexes, the IRIS-predicted top-1 pose corresponded to one of the lowest-ranked poses by RMSD, with deviations of 97–99 ranks from the true top-1 pose. Left: DNA duplex bound to 4′-hydroxymethyl-4,5′,8-trimethylpsoralen (1FHY). Middle: DNA duplex bound to spermine (1M69_2). Right: RNA riboswitch bound to 7,8-dihydroneopterin (7WII). Despite chemical diversity, these ligands share structural features that complicate pose re-ranking: either conformational flexibility (spermine) or planar heteroaromatics capable of multiple stacking orientations (psoralen and pterin). In each case, the binding environment (DNA grooves or a relatively shallow RNA pocket) provides limited geometric constraints, leading to multiple plausible docking configurations and reduced IRIS performance.

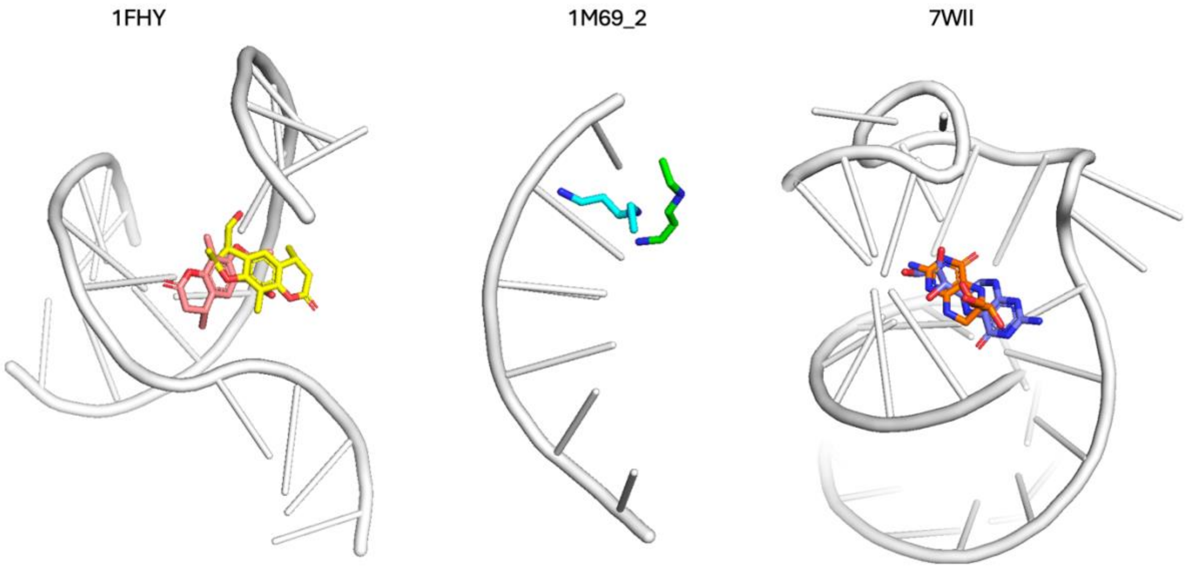

**Figure S4**

**Figure S4.** Normalized performance of IRIS trained on all nucleic acid complexes (teal) and IRIS trained exclusively on RNA complexes (coral) when evaluated on the RNA subset of the IRIS test set. Performance is expressed as the percentage of RNA test set complexes in which the Top Ranked pose (ranked #1 by each model) or the Best of Top 5 poses (lowest RMSD pose among the top five ranked poses) achieved an RMSD ≤ 2.0 Å. All values are normalized to the maximum achievable success rate for the RNA subset (82.41%), which reflects the percentage of RNA test set complexes for which rDock generated at least one of the 100 poses within 2.0 Å RMSD of the experimental structure

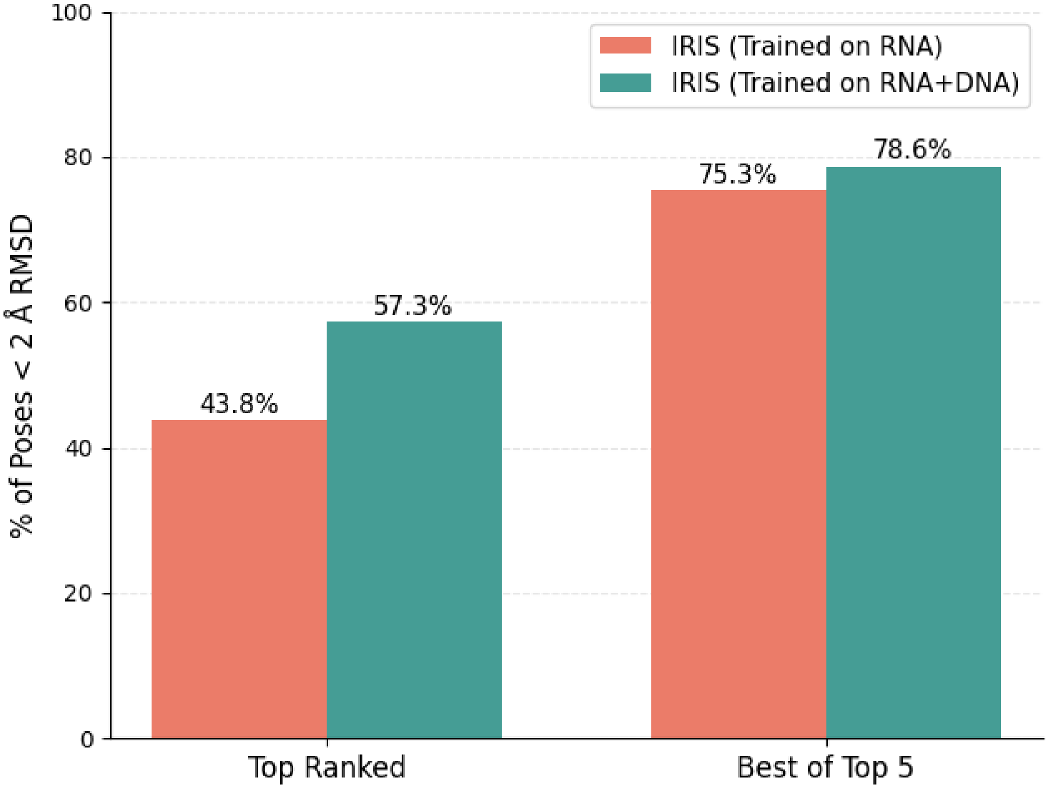

**Figure S5**

**Figure S5.** Relative SHAP contribution (%) of grouped feature categories to RMSD predictions for the RL_dock IRIS model. Bars indicate the summed mean absolute SHAP values for all features within a category, expressed as a percentage of the total summed mean absolute SHAP values across all features.

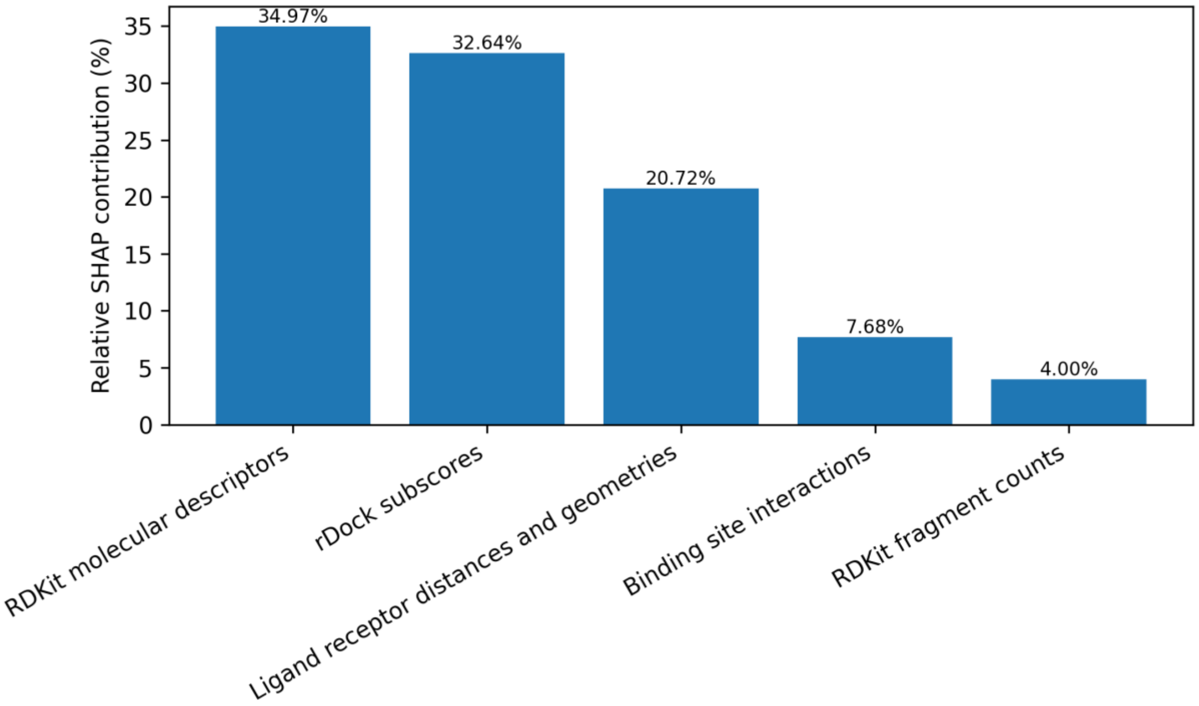

**Figure S6**

**Figure S6.** Learning curve for the IRIS RL_dock model: mean absolute error (Å) of predicted RMSD values relative to the true RMSD for ligand poses of complexes in the validation and training subsets as a function of training set size. Solid lines represent mean MAE values across 5 cross-validation folds and the shaded regions correspond to one standard deviation above and below the mean, calculated as $\mu \pm \sigma$, where $\mu$ is the MAE at each training set size and $\sigma$ is the standard deviation of the MAE across the cross-validation folds.

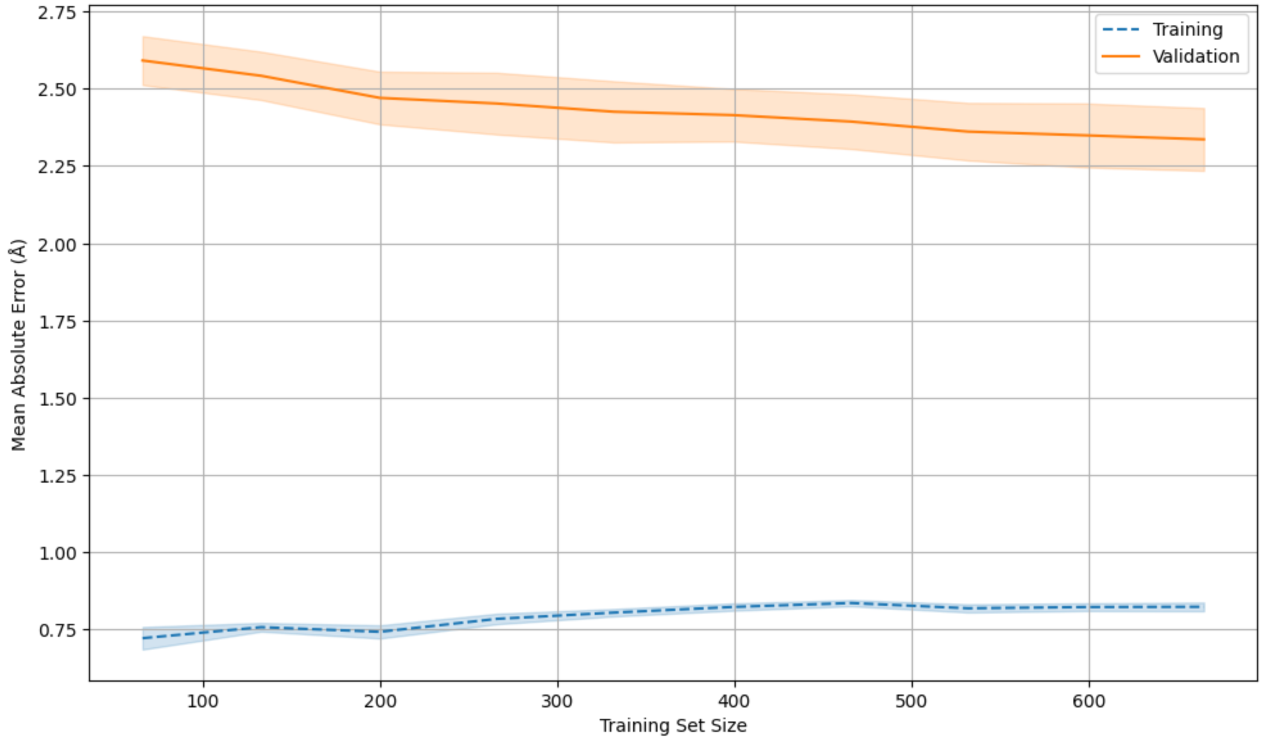

**Figure S7**

**Figure S7.** Top three complexes with the highest per-complex MAE values in RMSD prediction. From left to right: 1K2Z (MAE = 4.83 Å, Stdev = 1.24 Å), 3EUI (MAE = 4.84 Å, Stdev = 0.97 Å), and 453D ( MAE = 5.35 Å, Stdev = 0.44 Å), with ligands Distamycin A, 3-[(2R)-2-ethylpiperidin-1-yl]-N-[6-({3-[(2S)-2-ethylpiperidin-1-yl]propanoyl}amino)acridin-3-yl]propenamide, and 4-{[4-hydroxy-phenyl]-1h-benzimidazole-5-yl}-benzimidazole-2-yl-[4-hydroxy-benzene] respectively.

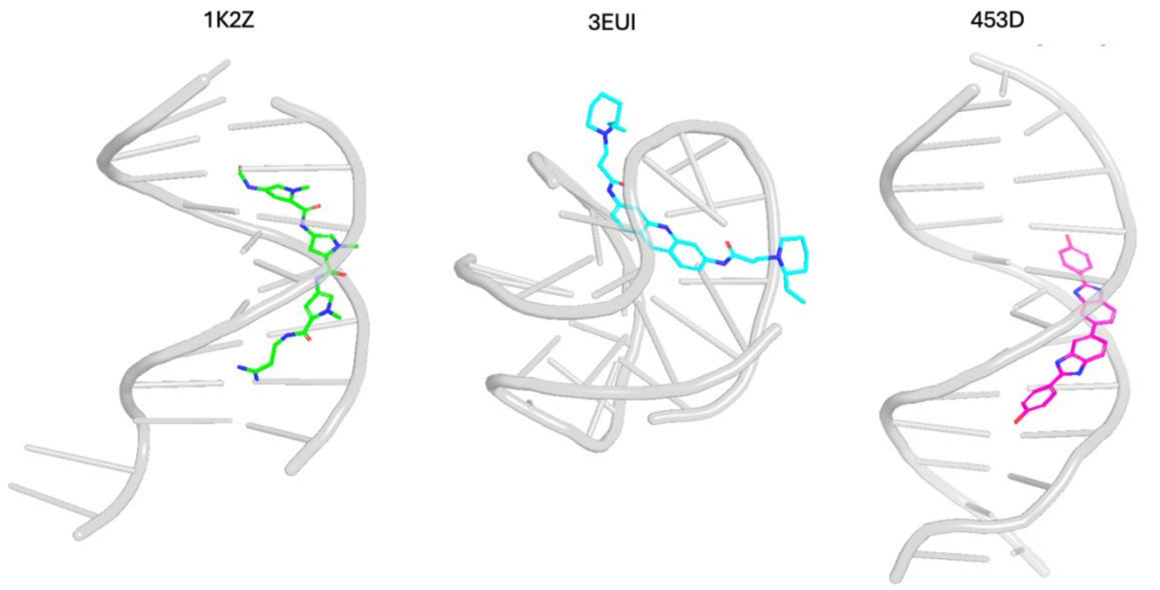

### Figure S8

**Figure S8.** Top 3 lowest MAE complexes. These complexes 2IBK (MAE = 0.61 Å ± 0.45 Å), 5BJP (MAE = 0.54 Å ± 0.29 Å), and 6D8A (MAE = 0.61 Å ± 0.33 Å), with ligands ethylene glycol, Dimethyl sulfoxide, and (4S)-2-Methyl-2,4-pentanediol respectively.

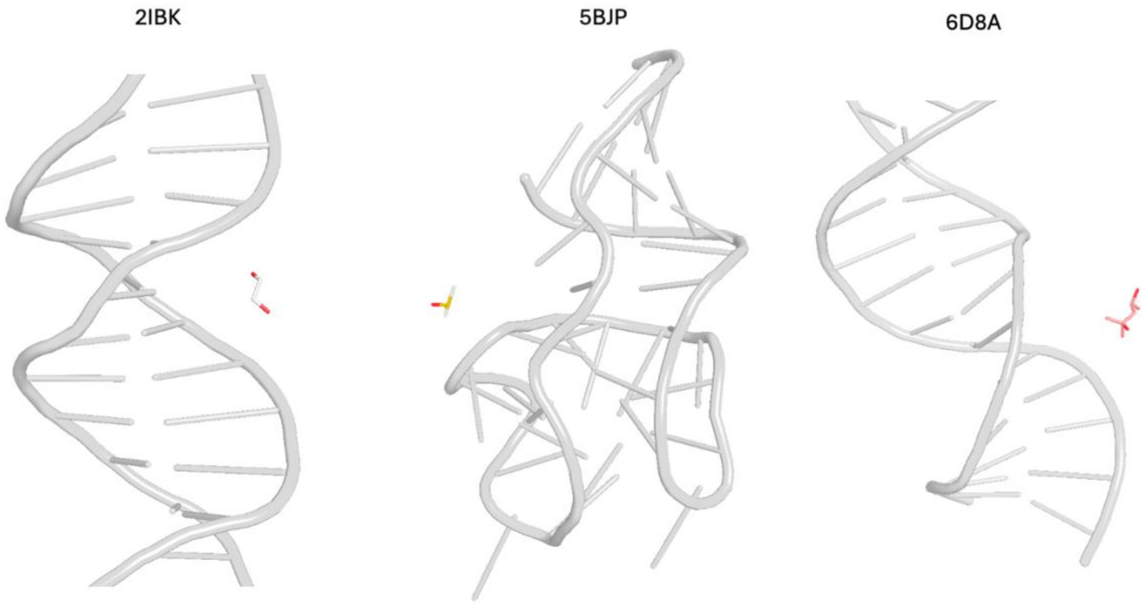

**rDock Docking Parameters**

Docking parameters used for the rDock local docking method, i.e., the Reference Ligand method, were set to generate 100 poses for each complex and configured according to the specified value ranges in the rDock reference guide35. The specific parameters are as follows: RECEPTOR_FLEX 3.0, SITE_MAPPER RbtLigandSiteMapper, RADIUS 4.0, SMALL_SPHERE 1.0, MIN_VOLUME 100, MAX_CAVITIES 1, VOL_INCR 0.0, GRIDSTEP 0.5.

Docking parameters used for the rDock global docking method, i.e., the Two Sphere method, were set to generate 100 poses for each complex and configured according to the specified value ranges in the rDock reference guide35. The specific parameters are as follows: RECEPTOR_FLEX 3.0, SITE_MAPPER RbtSphereSiteMapper, CENTER (x,y,z), RADIUS 12.0, SMALL_SPHERE 1.0, LARGE_SPHERE 6.0, MIN_VOLUME 100, MAX_CAVITIES 1. Here “(x,y,z)” denotes the coordinates of the center of mass of the native ligand
